# Spatially Resolved Phosphoproteomics Reveals Fibroblast Growth Factor Receptor Recycling-driven Regulation of Autophagy and Survival

**DOI:** 10.1101/2021.12.04.471203

**Authors:** Joanne Watson, Harriet R. Ferguson, Rosie M. Brady, Jennifer Ferguson, Paul Fullwood, Hanyi Mo, Katherine H. Bexley, David Knight, Jean-Marc Schwartz, Michael P. Smith, Chiara Francavilla

## Abstract

Receptor Tyrosine Kinase (RTK) endocytosis-dependent signalling drives cell proliferation and motility during development and adult homeostasis, but is dysregulated in diseases, including cancer. The recruitment of RTK signalling partners during endocytosis, specifically during recycling to the plasma membrane, is still unknown. Focusing on Fibroblast Growth Factor Receptor 2b (FGFR2b) recycling, we revealed FGFR signalling partners proximal to recycling endosomes (REs) by developing a Spatially Resolved Phosphoproteomics (SRP) approach based on APEX2-driven biotinylation followed by phosphopeptide enrichment. Combining this with traditional phosphoproteomics, bioinformatics, and targeted assays, we uncovered that FGFR2b stimulated by its recycling ligand FGF10 activates mTOR-dependent signalling and ULK1 at the REs, leading to autophagy suppression and cell survival. This adds to the growing importance of RTK recycling in orchestrating cell fate and suggests a therapeutically targetable vulnerability in ligand-responsive cancer cells. Integrating SRP with other systems biology approaches provides a powerful tool to spatially resolve celllar signalling.

## Introduction

Endocytosis is the process by which surface molecules, including Receptor Tyrosine Kinases (RTKs), undergo internalization from the plasma membrane (PM) into the early endosome (EEs) within seconds of ligand binding, followed by direct recycling to the PM, sorting to the lysosome via the late endosome (LEs) for degradation, or sorting to the recycling endosome (REs) for recycling to the PM^1–3^. In addition to controlling receptor availability at the cell surface^4^, recycling is critical for regulating signalling duration and output^1, 5–10^. For instance, we and others have established a link between the recycling of RTKs, such as Fibroblast and Epidermal Growth Factor Receptors (FGFR, EGFR) or the recycling of integrins and the sustained signalling activation that regulates cell motility^11–13^. Thus, recycling and its known regulators (e.g. RAB11, TTP, RCP)^1, 8, 12, 13^ maintain homeostasis in health and their dysregulation leads to several diseases, including cancer, diabetes, viral infections, and neurodegeneration ^8, 14, 15^. However, we still do not know which RTK signalling partners are recruited to the REs during recycling and thereby how they modulate downstream cellular responses.

The view of endocytosis as a way to attenuate signalling by receptor down-modulation and by controlling receptor availability at the cell surface has been challenged by data linking endocytosis to the propagation of RTK signalling from endosomes ^6, 7, 9^. For instance, EGFR signalling from EEs leads to AKT phosphorylation and cell survival ^16^, suggesting that EGFR internalization is required for the full spectrum of signalling activation downstream of EGFR. The recruitment of specific scaffold proteins to EEs regulates a certain branch of signalling, exemplified by the p14/MP1 complex engaging the kinase ERK-MAPK ^17, 18^. Another example of the crucial role of endosomes as signalling regulators is the recruitment of the LAMTOR complex to the lysosomes, which regulates the kinase mTOR in response to nutrients and growth factors with consequences for signalling, cell growth, metabolism, and autophagy ^19^. However, much less is known about the role of REs as signalling platform compared to EEs or lysosomes ^20, 21^. This knowledege would allow for specifically modulating cellular signalling. For instance, depleting the recycling adaptor RCP in cancer cells not only switches EGFR trafficking from recycling to degradation but also decreases cell proliferation and migration ^12^. Recently, we found a reciprocal regulation between FGFR2b and EGFR signalling outputs which i) occurs at the REs; ii) leads to FGFR2b-dependent phosphorylation of EGFR on threonine 696 (T693) and of the cell cycle regulator CDK1 on T161; iii) regulates cell cycle progression ^22^. This data suggests that the REs can integrate and propagate signals, prompting us to further investigate which FGFR2b signalling partners are specifically recruited at the REs.

The FGFR family is a useful model for studying the contribution of trafficking to signalling outputs^23^. There are four FGFRs, with FGFR1-3 having splice-variants denoted as b and c isoforms, and 21 FGF ligands, with each FGFR/FGF pair regulating signalling specificity in a context-dependent manner during development, in maintaining adult homeostasis, and in several diseases such as cancer ^24, 25^. One stark example of such functional selectivity is given by FGFR2b which is expressed on epithelial cells^24–26^. Stimulation of FGFR2b with FGF7 induced receptor degradation in contrast to stimulation with FGF10 which resulted in recycling of FGFR2b via the REs^13, 22, 27^. These two different trafficking routes of FGFR2b were associated with different phosphorylation dynamics within the signalling cascade and an increase in cell proliferation and proliferation/migration, respectively^13, 22^. Therefore, the duration and location of FGFR signalling must be strictly regulated to modulate the appropriate cellular outputs^23, 25^.

Here, to investigate FGFR2b signalling partners at the REs, we developed a Mass Spectrometry (MS)-based phosphoproteomics approach which allowed us to distinguish sites globally phosphorylated upon FGF10 binding to FGFR2b from those sites specifaclly phosphorylated in the proximity of the REs during receptor recycling. This Spatially Resolved Phosphoproteomics (SRP) approach is based on proximity-dependent biotinylation, which has been recently developed to profile the interactome of internalized receptors such as G-protein coupled receptors (GPCRs) and of stress granules^28–30^. Proximity-dependent biotinylation occurs when a bait protein, tagged with a biotin ligase such as BioID or a peroxidase such as APEX2, encounters other proteins within the labelling radius of 10 nm or 20 nm, respectively^31, 32^. Combined with biotin enrichment using streptavidin beads and MS analysis, interactomes of bait proteins can be identified for any subcellular compartment^33, 34^. Using peroxidases like APEX2 is preferable for investigating short-acting, dynamic processes, due to their short labelling time of 1 minute. This approach was successfully used to define the interactors of GPCRs and of selective autophagy receptor-dependent cargoes ^28, 35^ and was therefore our choice. However, we expanded the method by adding a phosphorylated peptide enrichment step *after* biotin enrichment of proteins using streptavidin beads and after protein digestion. This novel method has allowed us to uncover FGFR2b signalling partners localized at the REs and to study their impact on FGFR2b responses. To dissect the spatially restricted signalling modules regulated by FGF10/FGFR2b during recycling, we combined the SRP approach with traditional quantitative phosphoproteomics of epithelial cells where FGFR2b recycling was blocked^22^. We found novel FGFR2b signalling partners localized at the REs during recycling that regulate mTOR signalling with functional consequences for autophagy and cell survival.

## Results

### Inhibiting FGFR2b trafficking alters the phosphoproteome of epithelial cells

To investigate changes in FGFR2b signalling during recycling we have previously analysed the phosphoproteomes of cells stimulated with the recycling ligand FGF10^13, 22^. Here, we examined the effect of FGFR2b trafficking impairment on the FGF10-stimulated phosphoproteome of the epithelial cell lines HeLa, stably expressing FGFR2b (HeLa_FGFR2b^ST^), and T47D, which express endogenous FGFR2b^13, 22^. The transient expression in more than 80% of the cells of Dynamin_K44A-eGFP (dominant negative Dynamin, DnDNM2) or eGFP-RAB11_S25N (dominant negative RAB11, DnRAB11) inhibited FGFR2b internalization and recycling to the PM respectively, in response to FGF10 stimulation for 40 min, a time point where FGFR2b localized in the REs in cells expressing wild-type e-GFP-RAB11 (wild-type RAB11, wtRAB11) (Fig. 1a-b)^13, 22^. As shown for FGFR1^36^, also FGFR2b co-localized with the marker of EEs, EEA1, and with DnRAB11 in cells expressing DnRAB11 and was not found at the PM after longer stimulation with FGF10 (Fig. 1a-b)^22^, suggesting that FGFR2b is trapped in EEA1/DnRAB11-positive vescicles under this experimental condition. Therefore, expressing DnDNM2 and DnRAB11 impair FGFR2b trafficking and will be used here to study recycling-dependent changes in FGFR2b signalling in response to FGF10. Immunoblot analysis of HeLa, T47D, and BT20 (another breast cancer cell line expressing endogenous FGFR2b^22^) in the same experimental conditions showed that impeding FGFR2b trafficking did not alter FGFR2b activation or the phosphorylation of ERK1/2 downstream of FGF10, (Fig. 1c-e), as recently reported for Epidermal Growth Factor-stimulated cells^37^. Next, we used Mass Spectrometry (MS)-based quantitative phosphoproteomics to comprehensively investigate changes in FGFR2b signalling when FGF10-dependent FGFR2b trafficking was impaired. We stimulated both HeLa cells expressing FGFR2b and either eGFP (as control, HeLa-FGFR2b GFP), DnRAB11 (HeLa-FGFR2b DnRAB11) or Dn-DNM2 (HeLa-FGFR2b DnDNM2) and T47D transiently expressing wtRAB11, DnRAB11, or DnDNM2 with FGF10 for 40 min and analysed the proteome and the phosphoproteome by MS (Fig. 2a and Supplementary Fig. 1-2, Supplementary Tables 1-4). Firstly, we checked that the transient expression of dominant negative proteins did not alter the cellular proteome using Pearson correlation, which was indeed high among all the experimental conditions (Supplementary Fig. 1a and 1j and Supplementary Tables 1 and 3). The quality of the 7620 and 8075 phosphorylated sites quantified in HeLa-FGFR2b and T47D, respectively was consistent with previous publications^22^ (Supplementary Fig. 1c-g, l-p and Supplementary Tables 2 and 4). As HeLa-FGFR2b and T47D expressed different levels of FGFR2b and their proteome and phosphoproteome did nor correlate (Supplementary Fig. 2), we first focused on the HeLa-FGFR2b datasets and then used the results to interrogate the T47D datasets. Principal component analysis (PCA) separated the MS runs based on the experimental conditions (Fig. 2b), suggesting that the location of FGFR2b during FGF10-dependent FGFR2b trafficking affects global signalling activation. To characterize this, we utilised Fuzzy c-means clustering of phosphorylated sites significantly dysregulated across the four conditions (ANOVA, *p*-value < 0.0001) and identified 11 clusters (Fig. 2c-d and Supplementary Table 2). We focused on phosphorylated sites either regulated by FGF10 but not affected by the expression of DnRAB11 or DnDNM2 (clusters 3 and 4), sites regulated by FGF10 but dysregulated by the expression of DnDNM2 (clusters 5 and 8), or sites regulated by FGF10 but dysregulated by the expression of both DnRAB11 and DnDNM2 (clusters 9 and 11), hereby defined as the membrane, the internalization, and the recycling response clusters, respectively (Fig. 2d). Next, we used the 11 clusters identified in the HeLa-FGFR2b phosphoproteome as a training dataset to identify clusters with corresponding patterns of regulation in the T47D phosphoproteome, identifying the three response clusters corresponding to the membrane, internalization-dependent and recycling-dependent signaling (Fig. 2d-e and Supplementary Table 3). We therefore concluded that inhibiting FGFR2b trafficking affects the regulation of global signalling pathways regardless of FGFR2b levels and of the overall proteome, indicating the fundamental role of recycling in regulating FGFR2b signalling. Indeed, over-representation analysis (ORA) of KEGG pathways identified mTOR signalling as a pathway enriched for proteins dysregulated in the recycling response cluster common to both HeLa-FGFR2b and T47D cells (Fig. 2f). However, we did not find any signalling pathways specifically enriched upon inhibition of FGFR2b internalization only (cells expressing DnDNM2 and stimulated with FGF10) (Fig. 2f). This suggests that the FGFR2b recycling route and not merely the presence of FGFR2b presence in the cytoplasm regulates FGFR2b signalling in epithelial cells. We next used our recently developed tool SubcellulaRVis (https://www.biorxiv.org/content/10.1101/2021.11.18.469118v1) to analyse the subcellular localization of proteins in the mTOR signalling pathway belonging to the recycling response cluster and found an enrichment for vesicles, the REs, and the late endosome in both cell lines (Fig. 2g). Altogether, this data confirms that recycling is crucial for FGFR2b signalling and identifies the mTOR pathway as a key downstream effector. However, this approach did not reveal which FGFR2b signalling partners were recruited to and specifically phosphorylated in the proximity of the REs during receptor recycling.

**Fig. 1.**
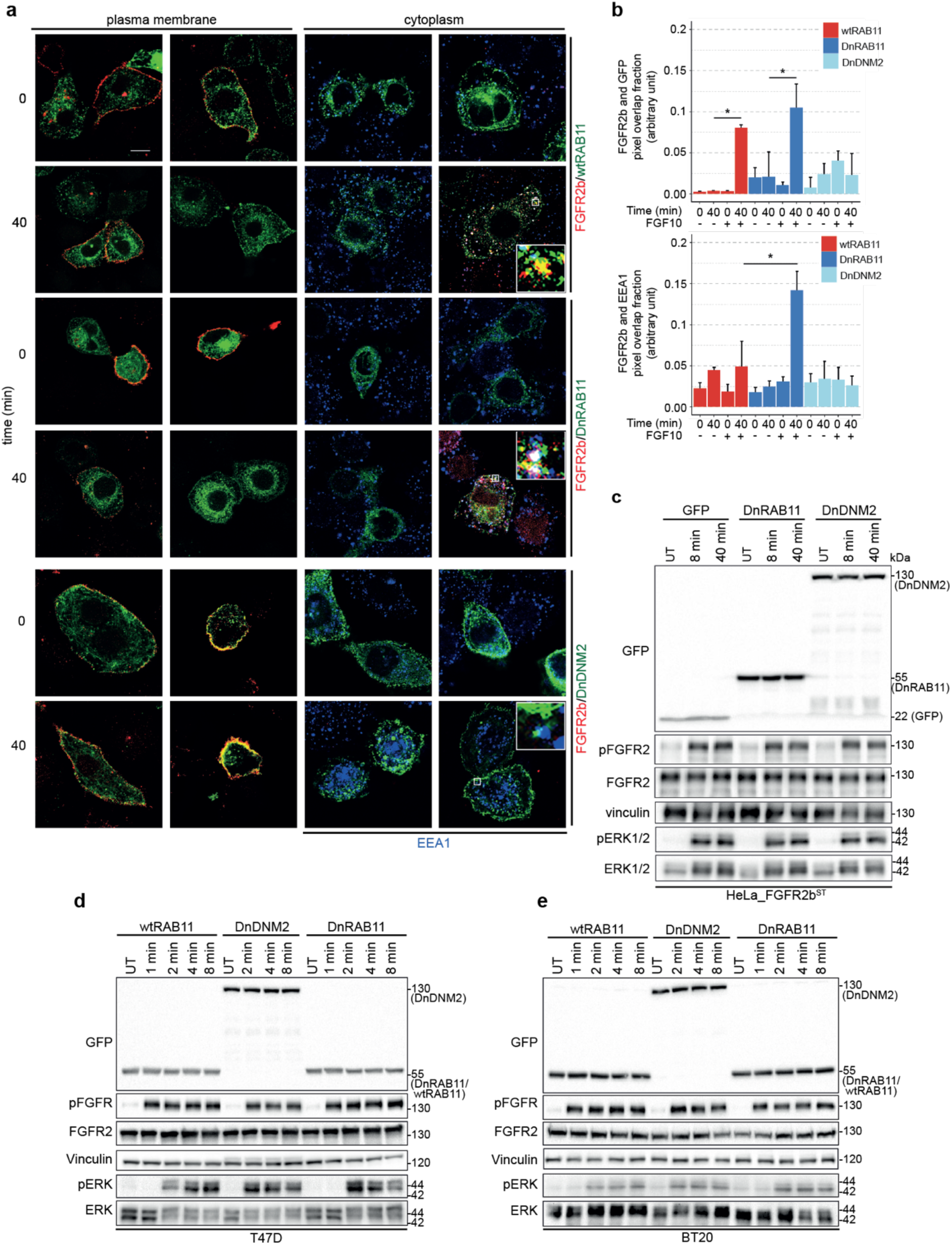
FGFR2b activation is not affected by receptor sub-cellular localization. **a** FGFR2b (red) internalization (cytoplasm) and FGFR2b recycling (plasma membrane) in HeLa cells stably transfected with FGFR2b-HA (HeLa_FGFR2b^ST^), expressing eGFP-RAB11a (wtRAB11), dominant negative eGFP-RAB11a_S25N (DnRAB11), or dominant negative dynamin-2_K44A-eGFP (DnDNM2) (green), and treated with FGF10 for 0 and 40 min. Early endosome antigen 1 (EEA1) (blue) is a marker for EEs^22^. Scale bar, 5μm. **b** Co-localization of FGFR2b (red pixels) with GFP-tagged proteins (green pixels) (top panel) indicated by red-green pixel overlap. Co-localization of FGFR2b (red pixels) with EEA1 (blue pixels) indicated by red-blue pixel overlap (bottom panel). Representative images are shown in 1A. Values represent median ± SD from N=3; * *p*-value < 0.005 (students t-test)^22^. Immunoblot analysis with the indicated antibodies of HeLa_FGFR2b^ST^ cells expressing GFP, DnRAB11 or DnDNM2 and left untreated (UT) or treated with FGF10 for 8 and 40 min (**c**), T47D (**d**) or BT20 (**e**) breast cancer cells expressing endogenous FGFR2b and transfected with wtRAB11, DnRAB11 or DnDNM2, and either left untreated (UT) or treated with FGF10 for 1, 2, 4 and 8 min.

**Fig. 2.**
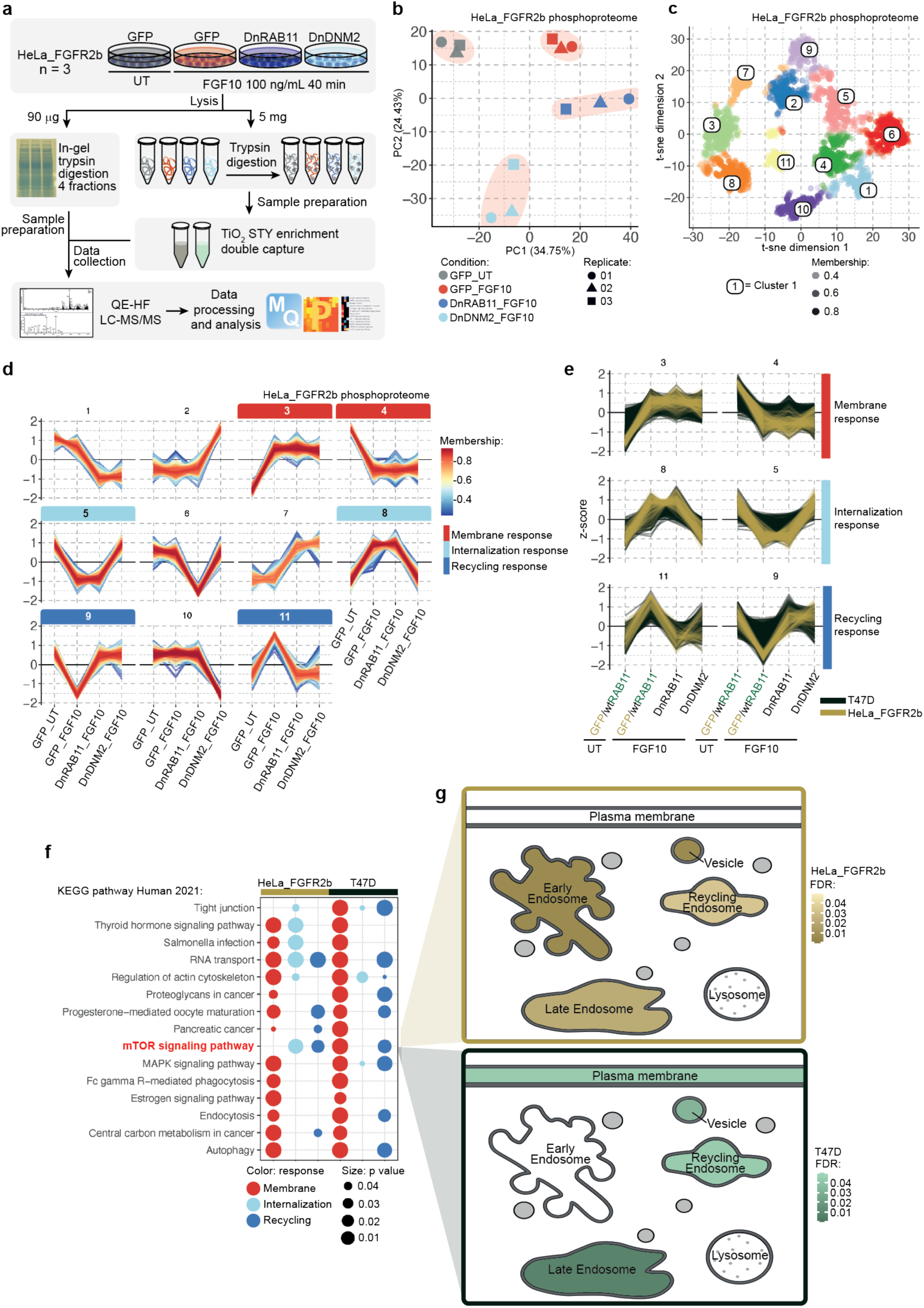
Phosphoproteomics analysis identifies FGFR2b internalization- and recycling-dependent signalling pathways. **a** Workflow of the phosphoproteomics experiment in HeLa cells transiently expressing FGFR2b (HeLa_FGFR2b) and either GFP, DnRAB11 or DnDNM2 and left untreated (UT) or treated with FGF10 for 40 min. **b** Principal Component Analysis (PCA) of the HeLa phosphoproteome from 2A showed small variation between technical replicates and separated samples based on experimental conditions. **c** t-distributed stochastic neighbour embedding (t-sne) analysis identified 11 clusters corresponding to phosphorylated peptides differentially regulated among four conditions. Colour corresponds to the cluster with highest membership score, determined using fuzzy c-means clustering based on the median z-score of the four conditions. Each cluster is identified by a unique colour and corresponding number. Opacity corresponds to the membership score assigned to each phosphorylated site within its most likely cluster. **d** Plots of the median z-scored intensities of phosphorylated sites based on the 11 clusters from Fig. 2c identified membrane response (red; clusters 3 and 4), internalization response (light blue; clusters 5 and 8), and recycling response (dark blue; clusters 9 and 11). Colour key indicates membership value assigned by Fuzzy c-means clustering. **e** Plots of the median z-scored intensities of phosphorylated sites based on the three main clusters identified in Fig. 2d. HeLa_FGFR2b (yellow) and T47D (dark green) were treated as in Fig. 2a and Supplementary Fig. 1h, respectively. **f** KEGG pathway over representation analysis (ORA) between HeLa_FGFR2b and T47D phosphorylated proteins within the membrane (red), internalization (light blue) and recycling (dark blue) responses (Fig. 2d) identified mTOR signalling as associated with FGFR2b recycling. The size of dot indicates statistical significance based on p-value. **g** Visualisation of the sub-cellular localization of the phosphorylated proteins belonging to the “mTOR signalling pathway” KEGG term in HeLa_FGFR2b (yellow) and T47D (green) using SubCellularVis (https://www.biorxiv.org/content/10.1101/2021.11.18.469118v1).

### APEX2-based proximity biotinylation assay enriches for phosphorylated proteins at the REs

To investigate recycling-dependent FGFR2b signalling in a spatially resolved (at the REs) and temporally sensitive (40 min simulation with FGF10) manner, we used the APEX2-based proximity labelling method ^33^ (Fig. 3a). The APEX2 method involves the fusion of a 27 kDa peroxidase enzyme to a bait protein (FGFR2b-HA, eGFP-wtRAB11, and eGFP in this study) that will rapidly biotin-label proteins within 20 nm of the bait protein in less than 1 min following addition of biotin-phenol (BP) and hydrogen peroxidase (H_2_O_2_) as an oxidant^32^. Biotinylated proteins can then be pulled-down using streptavidin beads and analysed using MS-based proteomics^33^. We designed an APEX2-based experiment to identify the signalling partners associated with FGF10-dependent FGFR2b that is localize at the RAB11-positive REs. We stably transfected HeLa or T47D cells with FGFR2b-HA-APEX2 (HeLa_FGFR2b-APEX2^ST^, T47D_FGFR2^KO^_FGFR2b-APEX2^ST^) and verified that FGFR2b signalling and trafficking were not altered by the presence of APEX2 upon FGF10 stimulation over time (Fig. 3b-d, Supplementary Fig. 3a). To exclude events occurring at the REs and in the cytoplasm independent of FGFR2b activation we expressed either eGFP-RAB11-APEX2 and eGFP-APEX2 in HeLa-FGFR2b^ST^ (HeLa-FGFR2b^ST^_RAB11-APEX2; HeLa-FGFR2b^ST^_GFP-APEX2), respectively (Fig. 3a). For all three APEX2-tagged bait proteins, biotin-phenol treatment for 40 min and H_2_O_2_ incubation for 1 min followed by streptavidin-beads pulldown (hereby pulldown) enriched for the bait protein, biotinylated proteins and also phosphorylated proteins (Fig. 3e and Supplementary Fig. 3b-c). Indeed, following 1 and 8 min FGF10 treatment, known interactors of FGFR2b, such as phosphorylated PLCγ and SHC, but not histone (H3), were identified in the pulldown without changes in their basal-level activation (Fig. 3e and Supplementary Fig. 3c). Next, to confirm that RAB11-APEX2 successfully enriched biotinylated proteins in the proximity of the REs during FGFR2b recycling, we immunoblotted the RAB11-APEX2 pulldown for RAB25 and HA-FGFR2b (Fig. 3f, g). RAB11-APEX2 also associated with other known markers of recycling, including RCP ^12^ (Supplementary Fig. 3d), confirming that our approach allows detection of proteins in the proximity of RAB11-positive REs. Taken together, this data supports the use of APEX2 to reveal phosphorylated signalling partners recruited to FGF10-stimulated FGFR2b at RAB11-positive REs.

**Fig. 3.**
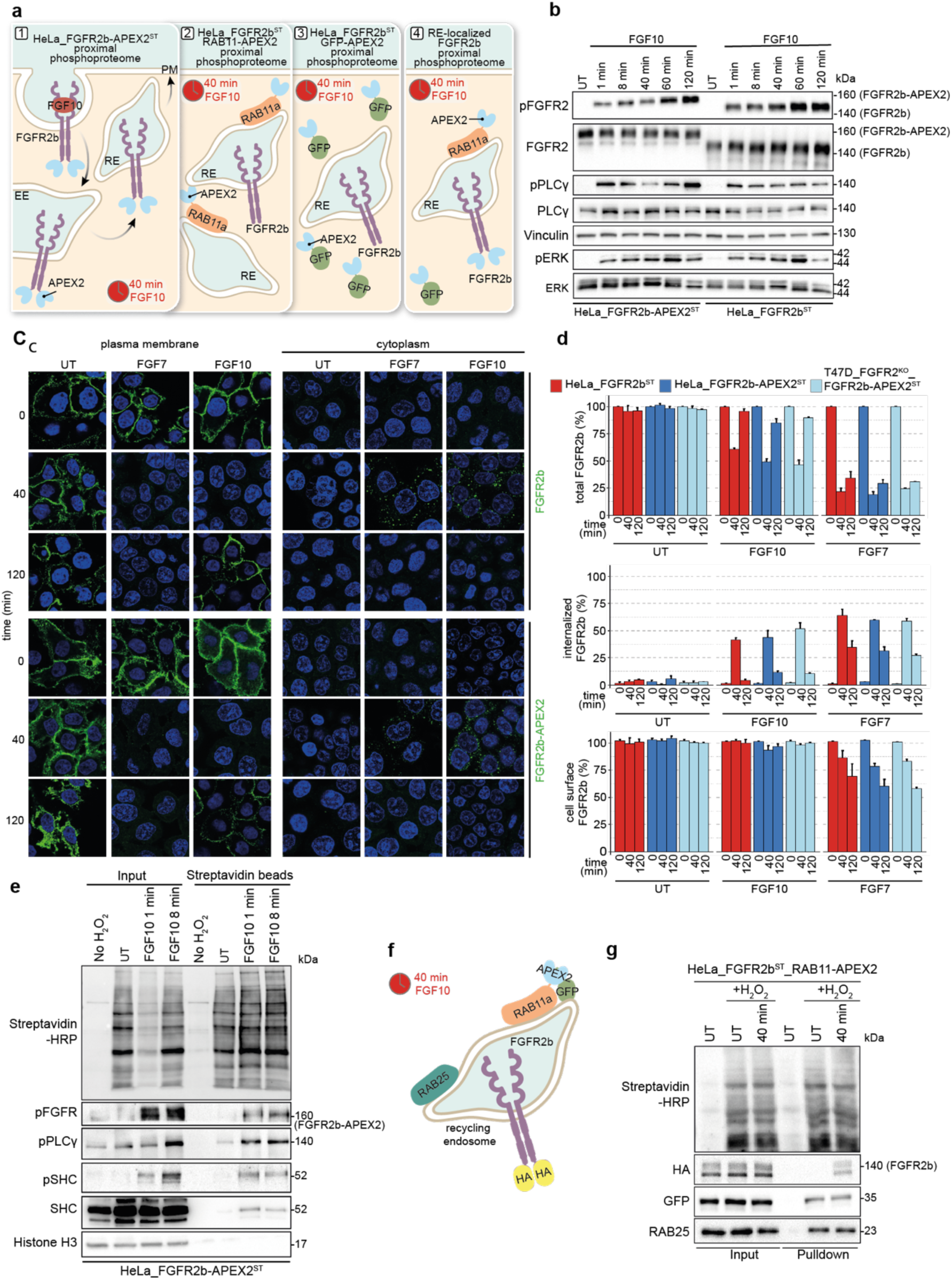
APEX2 tagged-FGFR2b and RAB11a identifies compartment-specific signalling partners upon FGF10 stimulation. **a** Schematic underlying the Spatially Resolved Phosphoproteomics (SRP) approach. Panel 1 represents the trafficking of FGFR2b-APEX2 stimulated with FGF10 in HeLa_FGFR2b-APEX2^ST^ and subsequent FGFR2b-APEX2 proximal phosphoproteome; panels 2 and 3 represent the localization of FGFR2b and of the APEX2-tagged proteins in cells expressing either FGFR2b and Rab11-APEX2 (HeLa_FGFR2b^ST^ RAB11-APEX2) or FGFR2b and GFP-APEX2 (HeLa_FGFR2b^ST^ GFP-APEX2) stimulated for 40 min with FGF10, and the proximal phosphoproteomes to the bait. Panel 4 represents the phosphorylated events occurring at the RAB11- and FGFR2b-positive REs upon 40 min FGF10 stimulation after subtracting cytosolic events using HeLa_FGFR2b^ST^ GFP-APEX2 proximal phosphoproteome. **b** Immunoblot analysis with the indicated antibodies of HeLa_FGFR2b^ST^ (right) or HeLa_FGFR2b-APEX2^ST^ (left) stimulated with FGF10 for 1, 8, 40, 60, or 120 min. **c** FGFR2b (top panels) and FGFR2b-APEX2 (bottom panels) (green) internalization (cytoplasm) and recycling (plasma membrane) in HeLa cells untreated (UT) or stimulated with FGF7 or FGF10 or 40 or 120 min. Nuclei are stained with DAPI (blue). Scale bar, 5μm. **d** Quantification of FGFR2b trafficking in HeLa_FGFR2b^ST^ (red), HeLa_FGFR2b-APEX2^ST^ (dark blue) and T47D_FGFR2^KO^_FGFR2b-APEX^ST^ (light blue) cells, showing the presence (total), internalized (internalized) and recycled (cell-surface) FGFR2b upon stimulation. Values represent mean ± standard error of mean (SEM) of at least three independent experiments. Representative images are shown in C. **e** Immunoblot analysis with the indicated antibodies of input or biotinylated proteins enriched with Streptavidin beads from HeLa_FGFR2b-APEX^ST^ left untreated (UT) and treated either with H_2_O_2_ or with FGF10 for 1 and 8 min. **f** Schematic of RE-localised FGFR2b, following 40 min of FGF10 treatment. Both RAB11-APEX2 and RAB25 localize at the REs^36^. **g** Immunoblot analysis with the indicated antibodies of input or biotinylated proteins enriched with Streptavidin beads from HeLa_FGFR2b^ST^_RAB11-APEX2 stimulated with either H_2_O_2_ or with FGF10 for 40 min.

### Spatially Resolved Phosphoproteomics (SRP) reveals FGFR2b signalling enriched at the REs

To uncover FGFR2b signalling partners at the REs in an unbiased manner, we designed a phosphoproteomics approach based on the detection of phosphorylated proteins in the pulldowns from cells expressing FGFR2b-APEX2 or RAB11-APEX2 (Fig. 3), hereby referred to as the Spatially Resolved Phosphoproteomics (SRP) approach (Fig. 4a). HeLa_FGFR2b-APEX2^ST^ expressing either RAB11, RAB11-APEX2 or GFP-APEX2 were treated with FGF10 for 40 min alongside HeLa_FGFR2b-APEX2^ST^ expressing RAB11 and HeLa-FGFR2b^ST^ GFP-APEX2 left untreated, as controls. We collected both the <u>global</u> proteome and phosphoproteome (obtained after TiO_2_-based chromatography enrichment of phosphorylated peptides) plus the <u>proximal</u> proteome and phosphoproteome obtained after enrichment of biotinylated proteins proximal to APEX2-tagged protein baits with streptavidin beads (proteome) followed by protein digestions and TiO_2_-based chromatography enrichment of phosphorylated peptides (phosphoproteome) (Fig. 4a and Supplementary Tables 5-6). We first analysed the global and the proximal proteomes, which both showed strong correlation between replicates and a clear distinction between the global and the proximal samples as assessed by Pearson correlation and PCA plots (Fig. 4b and Supplementary Fig. 4a-b). We noticed that the stimulated HeLa-FGFR2b^ST^ RAB11-APEX2 and the HeLa-FGFR2b^ST^ GFP-APEX2 proximal proteomes showed partial overlap in the PCA plots (Fig. 4b). To assess whether these were accurate spatial profiles of the REs and cytoplasmic background respectively, we checked which proteins were differentially abundant in each condition and performed an enrichment analysis for subcellular compartments on the resulting proteins lists (Fig. 4c-e). Proteins that were more abundant in the stimulated HeLa-FGFR2b^ST^ RAB11-APEX2 samples, including transferrin receptor (TFRC), VPS51 and VPS52^36, 38^ (Supplementary Tables 5-6), were enriched in the REs, whilst the proteins found in the stimulated HeLa-FGFR2b^ST^ GFP-APEX2 samples localized to the whole endosomal system (Fig. 4d-e). Therefore, our approach captured the spatial profile of REs at the proteome level and the prey proteins of the RE-specific bait protein RAB11-APEX2 can be differentiated from prey poteins of the GFP-APEX2 background control.

**Fig. 4.**
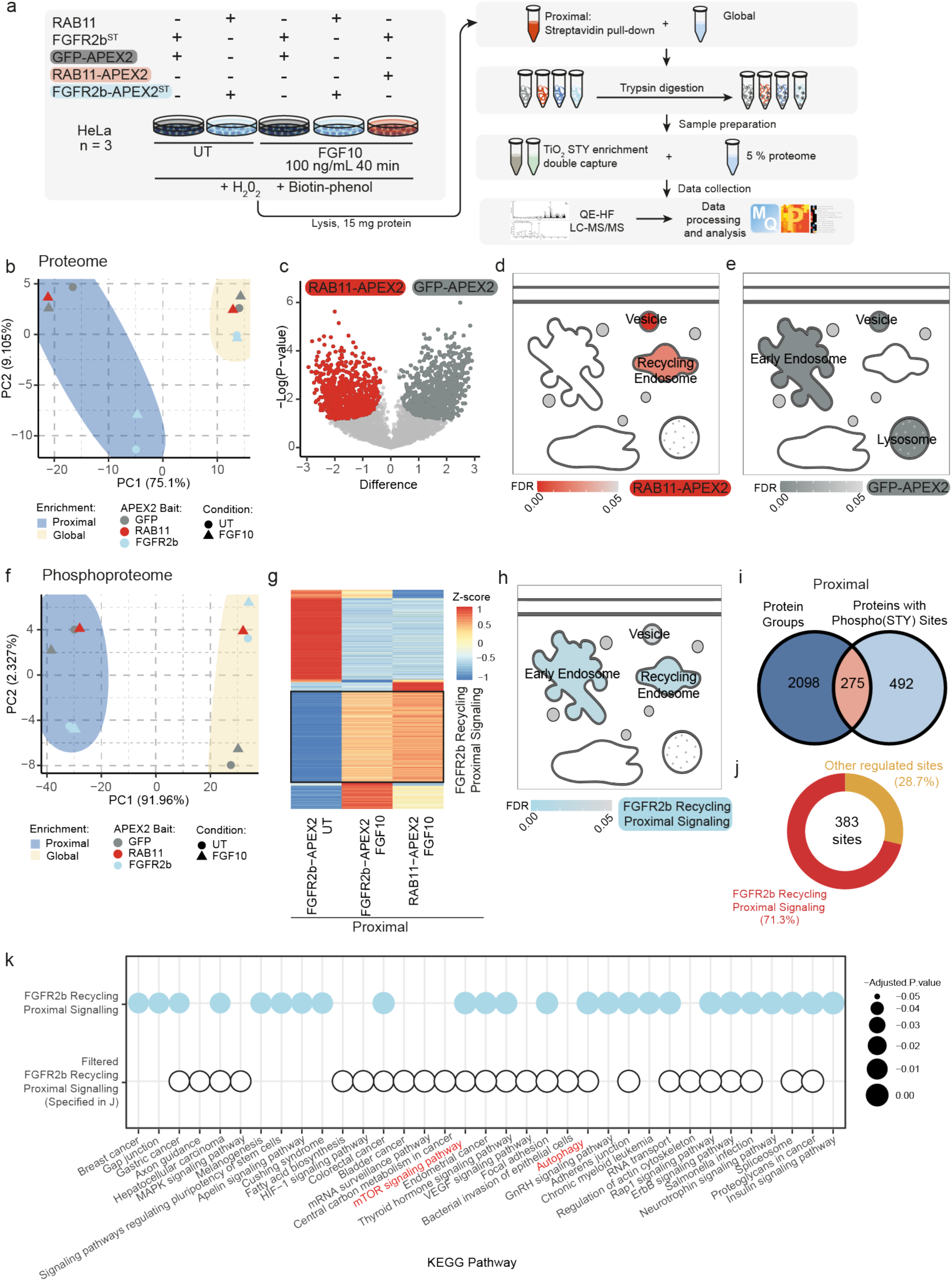
Spatially resolved proteomics and phosphoproteomics reveal FGFR2b-dependent regulation of mTOR signalling and autophagy. **a** Workflow of the spatially resolved proteomics and phosphoproteomics experiments in HeLa cells expressing the indicated constructs. **b** PCA plot of the proteome data based on median values. **c** Volcano plot of the proximal proteome comparing HeLa_FGFR2b^ST^ RAB11-APEX2 and HeLa_FGFR2b^ST^ GFP-APEX2 stimulated with FGF10 for 40 min. Visualisation of the sub-cellular localization of the differentially abundant proteins from HeLa_FGFR2b^ST^_RAB11-APEX2 (**d**) and HeLa_FGFR2b^ST^_GFP-APEX2 (**e**) stimulated with FGF10 for 40 min using SubCellularVis. **f** PCA plot of the phosphoproteome data based on median values. **g** Cluster analysis of the proximal phosphoproteome from the indicated conditions normalized to the proximal phosphoproteome of HeLa_FGFR2b^ST^ GFP-APEX2 for each timepoint. Phosphorylated sites upregulated at 40 min stimulation with FGF10 in both HeLa_FGFR2b^ST^ RAB11-APEX2 and HeLa_FGFR2b-APEX2^ST^ RAB11 are marked as the FGFR2b Recycling Proximal Signalling Cluster. **h** Visualisation of the sub-cellular localization of phosphorylated proteins found in the FGFR2b Recycling Proximal Signalling cluster using SubcellulaRVis (https://www.biorxiv.org/content/10.1101/2021.11.18.469118v1)**. i** Overlap of proteins and phosphorylated proteins detected in the proximal proteome and phosphoproteome samples, respectively. **j** Distribution of the 383 phosphorylated sites from 275 phosphorylated proteins, 71.3% of which were found in the FGFR2b Recycling Proximal Signalling Cluster. **k** KEGG pathway over representation analysis (ORA) of the phosphorylated sites found in the FGFR2b Recycling Proximal Signalling Cluster (blue light) and among the phosphorylated sites on proteins quantified at the proteome level from j (white).

Next, we analysed the phosphoproteomics data. Pearson correlation and PCA analysis revealed again strong correlation between replicates and a striking distinction between the global and the proximal samples (Fig. 4f and Supplementary Fig. 4c-d). Interestingly, the number of phosphorylated sites quantified in the proximal samples (2447) was substantially lower than the number quantified in the global samples (8545) (Supplementary Fig. 4e-f, Supplementary Table 6). This reduction was not seen in the number of quantified protein groups, indicating that the enrichment for phosphorylated peptides that follows the enrichment for biotinylated proteins had a substantial effect on the identification and quantification (Supplementary Fig. 4e-f). To assess how the double enrichment for biotinylated proteins followed by the enrichment for phosphorylated peptides affected the raw data analysis, we checked the confidence of identification and the distribution of intensities (Supplementary Fig. 4g-h). We found that there was a left-shift in the distribution of the intensities of the double-enriched proximal samples compared to the global samples, and therefore we normalized the global and the proximal phosphoproteome samples separately (Supplementary Fig. 4h). However, this analysis did not affect the overall quality of the proximal phosphoproteome data, as we found that >89% (6224 and 11726 for global and proximal, respectively) of the phosphorylated sites identified were Class I (≥ 0.75 localisation probability^39^) and had the expected proportions of single or multiple phosphorylated sites or serine, threonine, or tyrosine residues (Supplementary Fig. 4i)^22^. Finally, the APEX2 tag did not affect the quantification of the phosphoproteome (Supplementary Fig. 4j). We concluded that the double enrichment of biotynlated proteins and phosphorylated peptides does not impact data quality. To reveal the phosphorylated interactome of FGFR2b when localized at the REs we normalized the quantified phosphorylated sites from the control and from the FGF10-treated FGFR2b-APEX2 and RAB11-APEX2 samples against the corresponding time points of the GFP-APEX2 samples. Hierarchical clustering of the normalized data revealed a cluster of phosphorylated sites enriched in both the FGFR2b-APEX2 and the RAB11-APEX2 samples treated with FGF10, hereby the FGFR2b Recycling Proximal Signalling Cluster (Fig. 4g). This cluster was enriched for proteins localized to the EEs and the REs (representing the trafficking route taken by FGFR2b ^22^) (Fig. 4h) and included known regulators of trafficking like RCP, also known as RAB11FIP^12, 22^. Therefore, the stimulated proximal phosphoproteome captures phosphorylated proteins in the proximity of FGFR2b at the RAB11-positive REs. To confirm this, we analysed the overlap between phosphorylated proteins identified in the proximal phosphoproteome and proteins identified in the proximal proteome which would most likely represents phosphorylated FGFR2b partners at the REs. We found 275 proteins in common (Fig. 4i). The relatively small overlap (275 over 763 proteins with phosphorylated sites) may indicate the importance of performing the double enrichment step to reveal spatially resolved, phosphorylated signalling partners. Of the 383 phosphorylated sites on the 275 overlap proteins, 71.3% (273) were also found in the FGFR2b Recycling Proximal Signalling Cluster (Fig. 4j). Both the phosphorylated proteins belonging to the FGFR2b Recycling Proximal Signalling Cluster and the phosphorylated proteins found in the proximal proteome were enriched not only for pathways related to cancer and adhesion, but also signalling, including mTOR signalling, and for autophagy (Fig. 4k). These findings confirm that mTOR signalling is downstream of FGFR2b recycling (Fig. 2) and most likely activated when FGFR2b localizes at the REs and suggest a yet unexplored link between FGFR2b recycling, mTOR signalling and autophagy. Furthermore, these data indicate the robustness and specificity of the SRP approach to identify spatially resolved signalling pathways downstream of FGFR2b recycling^23–25^.

To verify whether the link between FGFR2b recycling, mTOR signalling and autophagy identified by the SRP approach was also evident from the global phosphoproteome data, we performed Volcano plot analysis of untreated and FGF10-stimulated global phosphoproteomes. We found differentially regulated sites in the two conditions as expected^13, 22^ (Fig. 5a), including FGFR2 phosphorylated sites (Y586, S587,Y656, Y657; Supplementary Table 6). However, there was only a small overlap (69 phosphorylated sites) between the FGF10-regulated phosphorylated sites from the global phosphoproteome and the FGFR2b Recycling Proximal Signalling Cluster, indicating that the SRP approach capably distinguished the proximal phosphoproteome from the global phosphoproteome (Fig. 5b). Indeed, whereas enrichment analysis of the upregulated phosphorylated sites from the global phosphoproteome revealed proteins involved in canonical FGFR signalling pathways, such as PI3K-AKT signalling, FOXO signalling and RAS signalling, the terms autophagy and mTOR signalling were enriched specifically in the FGFR2b Recycling Proximal Signalling Cluster (Fig. 5c). The proximal cluster also included significantly more connected phosphorylated proteins (443 compared to 224 found in the global phosphoproteome), indicating that FGFR2b localised to the RE has a densely interconnected, complex interactome (Fig. 5 c-d). FGFR2 and EGFR were found phosphorylated in both the proximal and the global phosphoproteome (Fig. 5e). One of the catalytic sites of FGFR2 (Y656) ^24^ was also identified as part of the internalization response cluster (Fig. 2), corroborating the role of this site for FGFR2 trafficking^13^. Interestingly, T693 on EGFR was found phosphorylated only in the proximal phosphoproteome (Fig. 5e), consistent with its role in regulating FGFR2b recycling at the REs^22^. Therefore, the SRP approach selectively enriches for phosphorylation proteins localized at the REs which could not be detected in the global phosphoproteome.

**Fig. 5.**
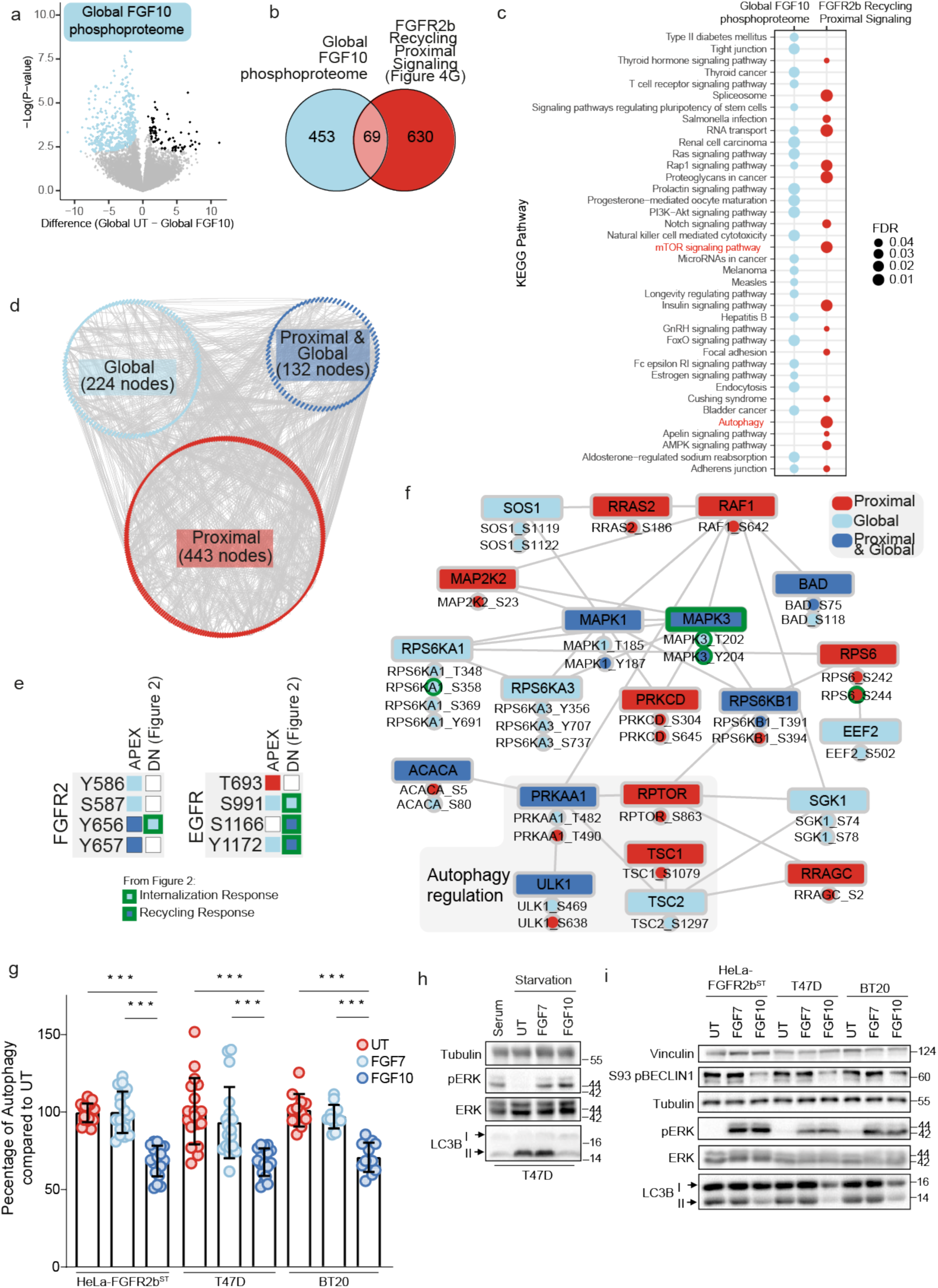
FGFR2b regulates mTOR signalling and autophagy from the REs. **a** Volcano plot of the phosphorylated sites from the global phosphoproteome in unstimulated and FGF10 40 min stimulated cells. **b** Overlap between the phosphorylated sites upregulated in the global phosphoproteome upon FGF10 stimulation based on A and the phosphorylated sites upregulated in the FGFR2b Recycling Proximal Signalling cluster from the proximal phosphoproteome (Fig. 4g). **c** KEGG pathway over representation analysis (ORA) of proteins with phosphorylated sites from b. **d** Protein-protein interaction network based on the STRING database and visualized with Cytoscape of the phosphorylated proteins shown in B. The number of phosphorylated protein nodes is indicated in parentheses. Unconnected nodes were removed. **e** Phosphorylated sites identified on FGFR2 and EGFR in the global (blue light) or in the proximal phosphoproteome (red) both (blue), and in the phosphoproteome from HeLa_FGFR2b^ST^ cells expressing GFP, GFP-DnRAB11 or GFP-DnDNM2 (Fig. 2a). Light blue with green border indicates phosphorylated sites found in internalization response clusters and dark blue with green border indicates sites found in recycling response clusters (Fig. 2d). **f** Subnetwork from D including proteins annotated to mTOR pathway or autophagy based on KEGG (c). Node colouring indicates whether the phosphorylated protein or the phosphorylated sites were found in global, proximal phosphoproteome or both. Proteins involved in autophagy regulation are highlighted in grey. Sites and proteins also quantified in Supplementary Table 2 have a green border. **g** Autophagy (measured by staining of lysosomes with acridine orange) of HeLa_FGFR2b^ST^, T47D, and BT20 untreated (UT) or treated with FGF7 or FGF10. N = 12, *p*-value =< 0.001*** (one-way ANOVA with Tukey test). **h** Immunoblot analysis with the indicated antibodies of the effect of serum starvation and FGF treatment on autophagic markers in T47D. LC3B II is the lapidated form. **i** Immunoblot analysis with the indicated antibodies of HeLa_FGFR2b^ST^, T47D, and BT20 treated with FGF7 or FGF10 for 2 h.

### FGFR2b recycling suppresses autophagy in a mTOR- and ULK1-dependent manner

Among the phosphorylated proteins annotated to the mTOR signalling pathway/autophagy in the enrichment analysis of the FGFR2b Recycling Proximal Signalling Cluster (Fig. 5c) we found several known components of mTOR signalling, including RAF1, RRAS2, MAP2K2, RPS6, and five proteins known to regulate autophagy either directly or via mTOR signalling: the AMPK subunit PRKAA1, the mTOR subunit RPTOR, TSC1, TSC2 and the kinase ULK1 phosphorylated on S638, the latter known to prevent autophagy^19, 40^ (Fig. 5f).

To test whether FGF10-mediated FGFR2b recycling regulates autophagy, we assessed autophagy firstly using acridine orange, widely used to stain lysosomes downstream of autophagy^41^, in HeLa-FGFR2b^ST^, T47D and BT20 treated for 2 h with FGF10 and with FGF7 (as a negative control for FGFR2b recycling^13^). FGF10 treatment significantly decreased autophagy compared to control in all cell lines, whereas FGF7 did not (Fig. 5g). As we starved cells before stimulation with FGFs and starvation is known to increase autophagy^42^, we checked the levels of known autophagy markers in starved cells followed or not by stimulation with either serum (as control), FGF7 and FGF10. The lapidated form (II) of the autophagosome-formation associated microtubule-associated proteins 1A/1B light chain 3B (LC3B)^43^ was supressed to levels seen in serum-treated cells by FGF10 treatment alone (Fig. 5h). This FGF10-, but not FGF7-dependent decrease in the levels of lipidayed LC3B (II) was seen in HeLa-FGFR2b^ST^ and BT20 cells as well, alongside a decrease in active BECLIN1 phosphorylated on S93 (Fig. 5i), another mediator of autophagosome formation and maturation^41, 44^. These results suggest that autophagy regulation is FGFR2b-recycling dependent downstream of FGF10. To confirm the importance of FGFR2b in autophagy regulation, we compared autophagy in parental T47D, T47D depleted of *Fgfr2*, and T47D depleted of *Fgfr2* and overexpressing FGFR2b (T47D_FGFR2b^KO^_FGFR2b^ST^) and found upregulation of autophagy in the absence of FGFR2b and less autophagy in T47D expressing high levels of FGFR2b compared to parental T47D (Supplementary Fig. 5a-b). Using the same cell model, we also investigated whether autophagy downstream of FGFR2b required mTOR signalling. To test this, we compared autophagy in cells subjected to either starvation, glucose removal and glucose-6-phosphate treatment (which are known to induce mTOR-dependent autophagy) or sodium valproate and fluspiriline treatment (which are known to induce mTOR-independent autophagy)^45, 46^. We found that mTOR-dependent but not mTOR-independent autophagy was affected by FGFR2b levels, as cells depleted of *Fgfr2* showed the highest levels of mTOR-dependent autophagy, whilst high levels of FGFR2b expression (T47D_FGFR2b^KO^_FGFR2b^ST^) induced the lowest levels of mTOR-dependent autophagy (Supplementary Fig. 5a-b). As high expression of RTKs may be associated with higher levels of internalisation in basal conditions and potential difference in signalling regulation ^47^ (Fig. 3c-d), we concluded that internalized FGFR2b was required for the regulation of mTOR-dependent autophagy. Moreover, mTOR-dependent, but not mTOR-independent, autophagy and the autophagy markers lapidated LC3B and BECLIN1 were regulated by FGF10 treatment in parental T47D (Supplementary Fig. 5c-d). Finally, both mTOR signalling and the known autophagy regulator ULK1 kinase were required for FGF10-depenent regulation of autophagy in T47D and HeLa cells (Fig. Supplementary Fig. 5e-f). These findings indicate that suppression of autophagy is FGFR2b recycling-, mTOR- and ULK1-dependent in FGF10-stimulated epithelial cells.

As we identified ULK1 phosphorylation on S638 in the proximal phosphoproteome (Fig. 5f) and this phosphorylation event is known to suppress autophagy^48^, we investigated whether phosphorylated ULK1 on S638 localized at the REs during FGFR2b recycling. In both HeLa-FGFR2b^ST^ and T47D cells ULK1 was phosphorylated on S638 in proximity of both FGFR2b and RAB11 (Fig. 6a-b and Supplementary Fig. 6a). Furthermore, confocal analysis of T47D_FGFR2^KO^_FGFR2b-APEX2^ST^ cells expressing wtRAB11 and stimulated with FGF10 for 40 min showed a significant co-localization between phosphorylated ULK1 on S638 and FGFR2b at the REs (Fig. 6c-d and Supplementary Fig. 6b). This findings confirm that ULK1 is associated to REs^49^ and suggest that the presence of stimulated FGFR2b at the REs is necessary for the recruitment of phosphorylated ULK1 on S638. Indeed, we did not visualize any ULK1 phosphorylated on S638 when FGFR2b recycling was impaired by expressing DnRAB11 (Fig. 6c-d). The phosphorylation of ULK1 downstream of FGFR2b recycling is a specific event, as other FGFR2b downstream pathways were only marginally affected in cells expressing either DnRAB11 or DnDNM2, treated with the primaquine and dynasore compounds, or stimulated with FGF7, all conditions that impaired FGFR2b trafficking^13, 22, 50, 51^ (Fig. 1, Fig. 6e-g and Supplementary Fig. 6c-d). Intriguingly, inhibiting FGFR2b localization at the REs also misplaced the FGFR2b recycling regulator TTP ^13^ from REs to LAMP1-positive lysosomes (Fig. 6c-d, h-i), where it has previously been shown to negatively regulate mTOR signalling^52^. Therefore, we checked mTOR localization and activation in our experimental conditions. mTOR was localized on lysosomes in both wtRAB11- and DnRAB11-expressing cells (Fig. 6c-d, h-i), but its activation decreased in cells with impaired FGFR2b trafficking (Supplementary Fig. 6e), confirming the link between FGFR2b recycling and mTOR signalling. Next, we checked whether inhibiting FGFR2b recycling by expressing DnRAB11 affected also mTOR signalling partners, including RAPTOR and AMPK^19^. Inhibiting FGFR2b recycling prevented RAPTOR phosphorylation and AMPK dephosphorylation on S863 and T172, respectively, events associated with increased mTORC1 activivty. Phosphorylation of S638 on ULK1 was also decreased up to 2 h after FGF10 stimulation when FGFR2b recycling was inhibited (Fig. 6g, Supplementary Fig. 6c).

**Fig. 6.**
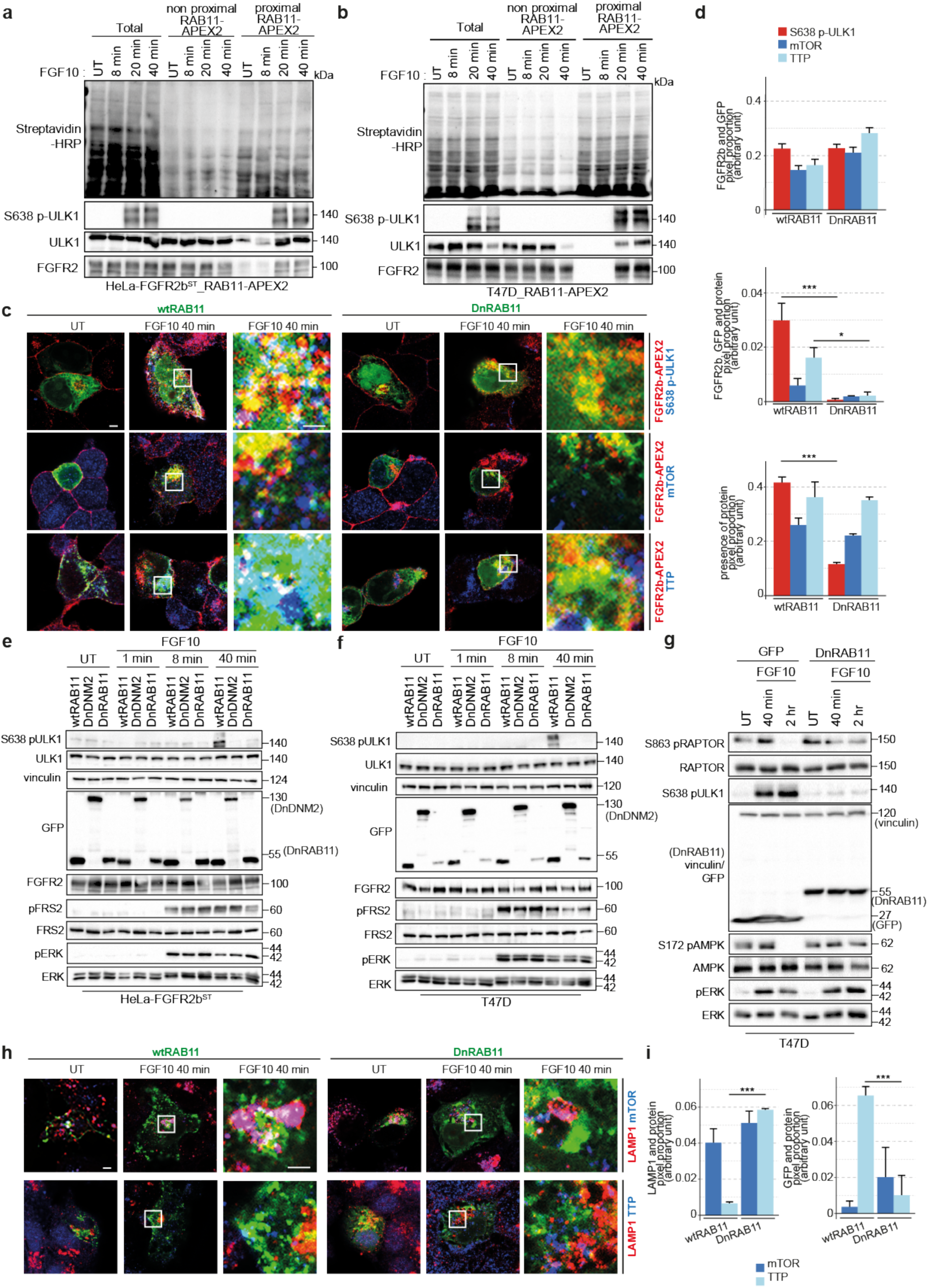
Phosphorylated ULK1 recruitment at the REs depends on FGFR2b recycling. **a, b** Immunoblot analysis with the indicated antibodies of HeLa_FGFR2b^ST^_RAB11-APEX2 (a) or T47D transfected with RAB11-APEX2 (T47D_RAB11-APEX2) (b) stimulated with FGF10 for the indicated timepoints. Non proximal and proximal samples represent the supernatant and the pulldown following enrichment of biotinylated samples with streptavidin beads, respectively, and run against total lysates (total). **c** Co-localization of FGFR2b-APEX2 (red) with phosphorylated ULK1 on S638, mTOR or TTP (blue) in T47D_FGFR2^KO^_FGFR2b-APEX^ST^ transfected with RAB11 or GFP-DnRAB11 (green) and stimulated or not with FGF10 for 40 min as indicated. Scale bar, 5 µm. The image on the right is a magnification of the FGF10-stimulated samples, scale bar 50 µm. **d** Quantification of pixels overlap of the conditions shown as representative images in C (see colour code) and of the proportion of each protein present in the indicated conditions based on pixels^22^. Values represent the median ± SD of at least three independent experiments. *p*-value < 0.005 **; *p*-value < 0.0005 *** (Students t-test). **e, f, g.** Immunoblot analysis with the indicated antibodies of HeLa_FGFR2b^ST^ (e) or T47D (f, g) transfected either with wtRAB11, DnRAB11, or DnDNM2 (e, f) or with GFP and DnRAB11 (G) and left either untreated (UT) or treated with FGF10 for the indicated time points. **h** Co-localization of LAMP1 (red) with mTOR or TTP (blue) in T47D_FGFR2^KO^_FGFR2b-APEX^ST^ transfected with wtRAB11 or DnRAB11 (green) and stimulated or not with FGF10 for 40 min as indicated. Scale bar, 5 µm. The image on the right is a magnification of the FGF10-stimulted samples, scale bar 50 µm**. i** Quantification of pixel overlap of conditions shown as representative images in H, Values represent the median ± SD of at least three independent experiments. *** *p*-value < 0.0005 (Students t-test).

In conclusion, FGFR2b recycling regulates mTOR signalling and the localization of phosphorylated ULK1 at the REs, with these signalling events being crucial for autophagy suppression downstream of FGF10.

### FGFR2b recycling regulates mTOR- and ULK1-dependent cell survival

To Investigate how FGFR2b signalling partners at the REs (e.g. ULK1) affected long-term FGFR2b responses during recycling, we tested the impact of impaired FGFR2b trafficking on FGF10-regulated responses. Firstly, we found that autophagy did not change or was slightly increased in FGF10-stimulated cells expressing DnRAB11 or treated with the trafficking inhibitors monensin^36^ and dynasore^50^ and in cells stimulated with FGF7 for 2 h (Fig. 7a and Supplementary Fig. 7a). Immunoblot analysis showed reduced levels of lapidated LC3B and RAPTOR phosphorylation on S863 and increased phosphorylation of ULK1 on S638 2 h following stimulation with FGF10 in wild type cells but not in cells expressing DnRAB11 or DnDNM2 (Fig. 7b and Supplementary Fig. 7b). Consistent with previous data, FGF7 did not affect any of these autophagy markers (Fig. 7b, Fig. 5g-i, Fig. 6e-g, Supplementary Fig. 6c-d and Supplementary Fig. 7b). Therefore, autophagy is regulated downstream of FGFR2b recycling in response to FGF10. Next, we treated cells with inhibitors of ULK1 (ULK101, SBI0206965) and of FGFR (PD173074) or mTOR (rapamycin) as controls, and investigated autophagy, apoptosis and proliferation following 24 h stimulation with FGF10 (Fig. 7c). ULK1 inhibition reduced autophagy compared to untreated cells but the level of autophagy was comparable to that of FGF10-treated cells, consistent with our data after 2h stimulation (Supplementary Fig. 5e-f). As expected, both mTOR and FGFR inhibitors increased autophagy and its markers in response to FGF10 (Fig. 7c, Supplementary Fig. 5e, f and Supplementary Fig. 7c). In addition, ULK and mTOR, but not FGFR, inhibition induced higher levels of apoptosis and of the apoptotic marker caspase 3 than in FGF10-treated cells (Fig. 7c and Supplementary Fig. 5c). All the inhibitors however induced a decrease in overall cell proliferation (Fig. 7c). Altogether, this data suggests a functional link between FGFR2b recycling, the activation of mTOR and the localization of phosphorylated ULK1 in proximity of the REs (Fig. 6) and long-term responses.

**Fig. 7.**
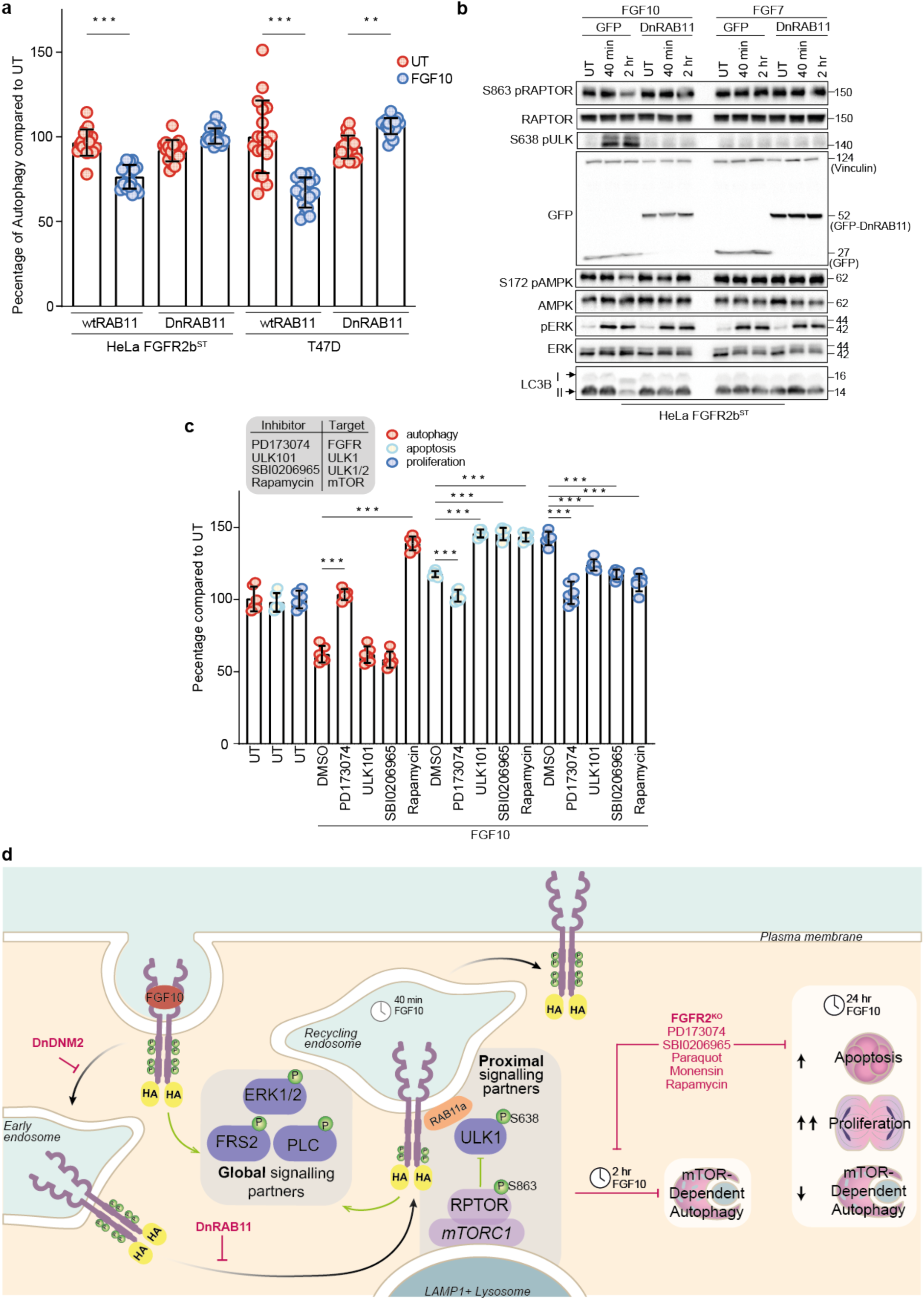
FGFR2b recycling regulates autophagy and the balance of proliferation and cell death. **a** Autophagy measured by acridine orange of HeLa_FGFR2b^ST^ (left) or T47D (right) transfected either with wtRAB11, DnRAB11, or DnDNM2 and incubated or not with FGF10 for 2h. N = 18, *p*-value =< 0.001*** (one-way ANOVA with Tukey test). **b** Immunoblot analysis with the indicated antibodies of HeLa_FGFR2b^ST^ transfected either with GFP or DnRAB11 and left either untreated (UT) or treated as indicated. **c** Measurement of cell proliferation by EdU incorporation, cell apoptosis by cleaved caspase 3 activated dye, and autophagy by acridine orange staining in T47D treated with the with PD173074, ULK101, SBI0206965, or Rapamycin which inhibit FGFR, ULK, ULK1/2, and mTOR (insert table) and stimulated or not with FGF10 for 2h. Data are presented as percentage compared to untreated cells. N = 6, *p*-value =< 0.001*** (one-way ANOVA with Tukey test). **d** Model of FGFR2b global and proximal signalling partners during recycling to the plasma membrane. Long-term responses are indicated based on the data of this study. The black arrow indicates FGFR2b trafficking. The green arrow indicates events activated by FGFR2b regardless of its subcellular localization.

In conclusion, we showed the importance of REs as signalling platforms to co-ordinate cellular fate by revealing that the inability of active FGFR2b to reach the REs disengages the link between mTOR/ULK1 signalling, autophagy and overall cell survival.

## Discussion

The importance of endocytosis in regulating selected RTK signalling cascades to drive cell fate in different contexts, including development or cancer, is now recognised ^8, 10, 14, 53, 54^. Here, we developed Spatially Resolved Phosphoproteomics (SRP) to uncover FGFR2b signalling partners localized at the REs during receptor recycling and we found mTOR-regulated players among them (Fig. 7d). We showed that the autophagy regulator ULK1 phosphorylated on S638 was recruited to FGFR2b- and RAB11-positive REs to prevent autophagy in FGF10-stimulated cells. The recruitment of phosphorylated ULK1 was prevented in the absence of FGFR2b recycling, resulting in impaired autophagy. Chemical inhibition of ULK1 and of mTOR, one of the known regulators of ULK1^48^, not only released FGF10/FGFR2b-dependet suppression of autophagy but also perturbed the longer-term effects on cell behaviour downstream of FGFR2b activation, including the balance between apoptosis and proliferation (Fig. 7d).

Inhibiting FGFR2b trafficking by genetic means alters the global phosphorylation programme in response to FGF10 (Fig. 2), confirming the crucial role of receptor internalization and recycling in driving signalling dynamics and long-term responses^5, 6, 13, 22^. Indeed, 24.56% and 13.6% of FGF10-dependent regulated phosphorylated sites depended on receptor internalization and recycling, respectively, in epithelial cells (Supplementary Table 2 and 4). This data highlights that certain signalling cascades are activated only when the receptor is “at the right place at the right time”^55^. Indeed, dysregulation of RTK trafficking leads to alteration in signalling activation such that endocytosis is now considered one of the hallmarks of health and diseases, including viral infections, neurodegeneration and cancer^8, 14, 15, 56^. For instance, the FGFR2b internalization-dependent phosphorylated sites discovered here could inform us on how the protein Dynamin regulates FGFR2b and, more broadly, RTK functions in breast cancer^57^. However, our genetic approach did not distinguish signalling partners specifically recruited to and phosphorylated at the REs during FGFR2b recycling. To reveal this, we developed a biotinylation-driven approach that we named Spatially Resolved Phosphoproteomics (SRP). This approach enabled us to generate a snapshot of spatially and temporally resolved signalling partners downstream of FGFR2b (Fig. 4 and Fig. 5), expanding the analysis from partners in the proximity of a protein bait as shown for GPCRs^28^ to *phosphorylated* partners in the proximity of FGFR2b at the REs. We first enriched for biotinylated proteins, and then for phosphorylated peptides (following protein digestion), a method already proven efficient with a BioID-based protocol^58^, but that differs from most of the published work in which enrichment for phosphorylated and APEX2-biotinylated proteins was performed prior to protein digestion^59^. Our method therefore resolves phosphorylated sites and not proteins, adding a layer of complexity in the analysis of cellular signalling. We envision that SRP could be easily adapted to study the localized dynamics of other post-translation modifications, thus enlarging the recently published BioID organelle interactome libraries^34^. The advantage of using APEX2-driven biotinylation over BioID or TurboID for defining signalling events in subcellular compartments is the tighter time frame that can be defined (e.g. seconds), which is essential for defining discrete signalling events^60^. The field of spatial proteomics is indeed growing, and novel technologies are in development to study spatially regulated cellular signalling^33, 61–63^. In contrast to other spatial phosphoproteomics methods (https://www.biorxiv.org/content/10.1101/2021.02.02.425898v1) that enrich for organelles and then analyse global phosphoproteomics, the SRP approach allows the enrichment of phosphorylated proteins at one (or more) organelle under acute stimulation, thus revealing unique phosphoproteome signatures in a spatio-temporal defined manner. In addition, our SRP method provides the exciting opportunity to investigate endosome-proximal phosphorylation events in a high-throughput manner as opposed to signalling partners identified at EEs using biochemical and low-throughput methods. For example, populations of phosphatidylinositol 3-kinase (PI3K) have been shown to be activated at the EEs downstream of EGF stimulation^64^. Signalling from the LEs has also been described, via MAPK and JNK^65^. However, signalling partners localized at RAB11-positive REs are less known. Here, SRP identifies 693 phosphorylated sites in proximity of the FGFR2b- and RAB11-positive REs, among which was ULK1 phosphorylated at S638.

Phosphorylated ULK1 at S638 is primarily seen at the REs when stimulated FGFR2b is also localized to REs. ULK1 has been previously shown to localize to RAB11-positive REs, but its phosphorylation state and the intersect with RTK trafficking have not been previously investigated^48, 49^. We also showed that upstream regulators of ULK1, such as mTOR or AMPK^48^, are in the proximity of the REs, suggesting that the REs may be a site of ULK1 regulation downstream of FGFR2b. The link between mTOR and endocytic trafficking processes such as lysosomal transport has been previously reported^66–68^. A potential dynamic interface between the REs and lysosomes, which are localized in close proximity one to each other^69^ would favour the interaction of mTOR complexes and ULK1 within the REs to drive downstream cellular responses. This intriguing possibility is worthy of further investigation – including the analysis of the role of AMPK, another ULK1 regulator^48^. This would have important implications for the understanding of how RTK trafficking and in general the endosomal system, regulates signalling specificty.

RTK signalling and endocytosis have previously been linked to regulation of autophagy^70^ and EGFR recycling has been shown to decrease in cells lacking autophagy regulators^71^. Signals from growth factors are known to converge on the mTORC1 complexes at the lysosomal membrane to inhibit autophagy and catabolic processes^19^. Focusing on the FGFR family, the FGFR2b selective ligand FGF7 has been shown to induce autophagy in keratinocytes after 24 h stimulation^72^ and FGF signalling regulates bone growth through autophagy^73^. However, within the 2 h timeframe used in our experiments, FGF7 fails to alter ULK1/mTOR signalling or the downstream autophagy response, in contrast to the responses achieved in FGF10-stimulated cells. The stark difference between FGF7 and FGF10 high lights the role of FGFR2b recycling as the regulator of the FGF10/ULK1/autophagy interplay. How this is orchestrated from the REs remains however unclear. One possibility is the involvement of EGFR signalling, as we have recently shown that EGFR is phosphorylated downstream of FGF10/FGFR2b recycling at the REs^22^ and EGFR signalling regulates autophagy^74^ with EGFR trafficking requiring autophagy regulators^71^. Alternatively, REs and autophagosomes share signalling regulatory components that would require further investigations^49, 75^. Thus, a picture of REs as a point of convergence for several signalling pathways and for coordination of long-term responses is clearly emerging. This information can be used to exploit REs for nanomedicine, for instance for a better deliver of siRNA against specific signalling players^76^.

Recycling is known to control cellular responses, including proliferation, migration, invasion and, as shown in this study, the rate of autophagy^11–13^. It is therefore not surprising that impeding the recycling of FGFR2b leads to dysregulated cellular proliferation and cell death with broader implications for the spatio-temporal regulation of FGFR signalling^77^. We have started dissecting how cell proliferation is tightly regulated by multiple converging mechanisms downstream of FGFR2b, including receptor recycling and its duration^13, 22^, EGFR, CDK1 and ULK1 phosphorylation occurring at the REs^22^ (Fig. 7), and suppression of autophagy (Fig. 7).

The G1/S checkpoint is known to be controlled by the homeostatic balances of nutrients, such as amino acids and sugars, all regulating mTOR signalling^78^. Thus, FGF10 may specifically alter the balance at this checkpoint by supressing the negative regulation of autophagy on cell cycle progression via mTOR/ULK1 during receptor recycling^78–80^. Alternatively, FGF signalling could regulate the link between cell cycle, number, and size by controlling the activation of CDKs, mTOR, and MAPKs, respectively^81^. The combination of pharmacological inhibition of signalling, autophagy, and mTOR signalling inhibitors has shown greater cytotoxic effects in several diseases^82^. It is therefore time to speculate that such a combination may prove efficient in FGFR2b-driven genetic diseases or cancer, including breast cancer^83, 84^.

In conclusion, we discovered a role for internalized FGFR2b in regulating autophagy from the REs. The approach described here and the datasets collected provide a resource to the cell signalling research community and can be used to further study the role of internalized, activated RTKs in modulating signalling cascades.

## Acknowledgements

We thank Professor Martin Lowe and Professor Alan Whitemarch, for reading the manuscript and the Bioimaging and the Bio-MS Facilities (University of Manchester). We thank Dr A. Badrock, Hurlstone, and Wilcock for reagents and all members of the Francavilla’s team for discussion. Research in CF lab is supported by the Wellcome Trust (107636/Z/15/Z and 107636/Z/15/A), the Biotechnology and Biological Sciences Research Council (BB/R015864/1), and Medical Research Council (MR/T016043/1). PhD students are supported by BBSRC Doctoral Training Programme (HF and JW: BB/M011208/1. For the purpose of open access, the author has applied a CC BY public copyright licence to any Author Accepted Manuscript version arising from this submission.

## Author contributions

MPS performed experiments, supervised HRF, RMB, and KHB, and wrote the manuscript. HRF and JW performed experiments and data analysis. RMB, KHB, and JF performed experiments. HM helped with data analysis. PF contributed to sample preparation for MS analysis. DK provided technical advice for the MS experiments. JMS supervised JW and HM. CF conceptualized the study, acquired funding, supervised the work, and wrote the manuscript. All the authors contributed to writing and approving the manuscript.

## METHODS

### Plasmids, antibodies and reagents

Plasmids: eGFP-RAB11 (Addgene #12674); eGFP-RAB11_S52N, mutagenesis of eGFP-Rab11^13^; eGFP (Addgene #34680); dynamin-2_K44A-eGFP, Mutagenesis of Dynamin-eGFP^13^; HA-FGFR2b^13^; APEX2 (Addgene #49386); pCDH-EF1-HA-FGFR2b-T2A-mApple, pCDH-EF1-HA-FGFR2b-APEX2-T2A-mApple, HA-FGFR2b-APEX2, eGFP-RAB11-APEX2, eGFP-APEX2 (Generated for this study).

Antibodies: purchased from Cell Signaling Technology: p44/42 MAPK (Erk1/2) (137F5) Rabbit mAb (#4695); FGF Receptor 2 (D4L2V) Rabbit mAb (#23328); GFP (D5.1) Rabbit mAb (#2956); Phospho-FGF Receptor (Tyr653/654) (55H2) Mouse mAb (#3476); Phospho-PLCγ1 (Tyr783) Antibody (rabbit polyclonal) (#2821); PLCγ1 Antibody (rabbit polyclonal) (#2822); Phospho-SHC (Tyr239/240) Anitbody (#2434S); Shc Antibody (rabbit polyclonal) (#2432); Phospho-FRS2-α (Tyr196) Antibody (rabbit polyclonal) (#3864); LC3B Antibody (rabbit polyclonal) (#2775); Phospho-Beclin-1 (Ser93) (D9A5G) Rabbit mAb (#14717); Beclin-1 (D40C5) Rabbit mAb (#3495); Phospho-ULK1 (Ser638) (D8K9O) Rabbit mAb (#14205); ULK1 (D8H5) Rabbit mAb (#8054); Raptor (24C12) Rabbit mAb (#2280); Phospho-AMPKα (Thr172) (40H9) Rabbit mAb (#2535); AMPKα (D5A2) Rabbit mAb (#5831); mTOR (7C10) Rabbit mAb (#2983); Rab7 Anitbody (#2094S); Cleaved Caspase-3 (Asp175) Antibody (rabbit polyclonal) (#9661S); purchased from Sigma-Aldrich: Monoclonal Anti-γ-Tubulin antibody (mouse) Monoclonal Anti-γ-Tubulin antibody produced in mouse (#T5326); Anti-Vinculin antibody, mouse monocolonal (#V9264); Anti-HA (12CA5 (mouse monoclonal); purchased from Abcam: Anti-Histone H3 antibody - Nuclear Marker and ChIP Grade (rabbit polyclonal) (#ab-1791); Anti-Rab25 antibody (rabbit polyclonal) (#ab45855); Anti-LAMP1 – Lyososome Marker (rabbit pol-yclonal); purchased from Invitrogen: Goat anti-Rabbit IgG (H + L) Secondary Antibody, Alexa Fluor® 488 (#A11034); Goat anti-Mouse IgG (H + L) Secondary Antibody, Alexa Fluor® 488 (#A11001); Goat anti-Rabbit IgG (H + L) Secondary Antibody, Alexa Fluor® 568 (#A11011); Donkey Anti-Mouse IgG (H + L) Secondary Antibody, Alexa Fluor® 647 (#A31571); Donkey Anti-Rabbit IgG (H + L) Secondary Antibody, Alexa Fluor® 647 (#A31573); purchased from other suppliers: Anti-ERK 1/2 Antibody (MK1) (mouse monoclonal) (Santa Cruz Biotechnology, #sc-135900); Anti-FRS2 Antibody (mouse monoclonal) (Santa Cruz Biotechnology, #sc-17841); Purified Mouse Anti-EEA1 (monoclonal) (BD Bioscience, #610457); Mouse LAMP-1/CD107a Lumenal Domain Antibody (polyclonal) (R&D systems, #AF4320); CHMP1b (rabbit polyclonal) (Proteintech, #14639-1-AP); FIP1/RCP Antibody (rabbit polyclonal) (Novus Biologicals, #NBP2-20033); Peroxidase-AffiniPure F(ab’)2 Fragment Goat Anti-Mouse IgG (H + L) (Stratech, #115-036-045); Peroxidase-AffiniPure F(ab’)2 Fragment Goat Anti-Rabbit IgG (H + L) (Stratech, #115-036-062).

## EXPERIMENTAL MODELS

### Cell Culture

Human epithelial cell lines were purchased from ATCC, authenticated through short tandem repeat (STA) analysis of 21 markers by Eurofins Genomics, checked monthly for mycoplasma via a PCR-based detection assay (Venor®GeM – Cambio), and grown in the indicated media supplemented with 2 mM L-glutamine and 100 U/ml penicillin, 100 μg/ml streptomycin and 10% fetal bovine serum. HeLa, BT20 and Lenti-XL cells were grown in StableCell™ DMEM - high glucose (Sigma-Aldrich). T47D were grown in RPMI 1640 Medium, GlutaMAX™ Supplement (Gibco).

### Transfection

All transfections were carried out in Gibco opti-MEM glutamax reduced serum media (ThermoFisher Scientific). HeLa cells were transfected using Lipofectamine 2000 (ThermoFisher Scientific) according to the manufacturer’s instructions, 24 h after RNA interference transfection where indicated. T47D and BT20 cells were transfected using Escort IV according to manufacturer instructions, same as above. Assays were performed 36 h after transfection, as previously described^22^.

HeLa cells stably expressing HA-FGFR2b or HA-FGFR2b-APEX2 are referred to as follows: HeLa_FGFR2b^ST^, HeLa_FGFR2b-APEX2^ST^. Cells were transiently transfected with the following constructs: eGFP (GFP in text), dynamin_K44A-eGFP (DnDNM2 in text), eGFP-RAB11_S52N (DnRAB11 in text), eGFP-RAB11 (wtRAB11 in text), eGFP-RAB11-APEX2 (RAB11-APEX2 in text), eGFP-APEX2 (GFP-APEX2 in text).

### T47D cells depleted of *Fgfr2*

Guide RNAs (crRNA) (IDT) specific to *Fgfr2* were combined with a common trans-activating crRNA (Alt-R^®^ CRISPR-Cas9 tracrRNA) (IDT) to create a functional ribonucleoprotein (RNP) duplex. These were then pre-complexed with Cas9 nuclease (Alt-R^®^ S.p. Cas9 Nuclease V3) (IDT) and transiently transfected into parental T47D using Viromer® CRISPR transfection reagent (Cambridge Bioscience) according to manufacturer’s instructions. Colonies were selected and screened for high frequencies of genomic editing using the free Inference of CRISPR Edits (ICE) analysis tool by Synthego. Loss of protein expression was then confirmed by Western blot (see Supplementary Fig. 2a). T47D depleted of Fgfr2 are referred to as T47D_FGFR2b^KO^.

**Table 1:**
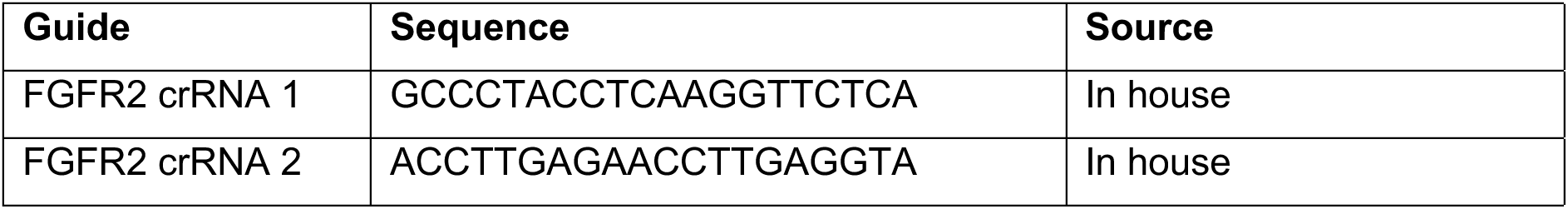
Guide RNAs for FGFR2 depletion

### Lentiviral transduction

Lenti-X cells were transfected in Gibco opti-MEM Glutamax reduced serum media (ThermoFisher Scientific) with the pCDH-EF1-HA-FGFR2b-T2A-mApple or pCDH-EF1-HA-FGFR2b-APEX2-T2A-mApple viral vectors, alongside VSV-G envelope expressing plasmid pMD2.G and lentiviral packaging plasmids pMDLg/pRRE and pRSV-Rev (all generously gifted from Dr Hurlstone), using FuGENE^®^ HD Transfection Reagent (Promega), following manufacturer instructions. After 48h, the lentivirus-containing media was sterile filtered using 0.22 μm syringe filter and stored at -80°C. The lentiviral-media was added to HeLa or T47D_FGFR2b^KO^ in Gibco opti-MEM Glutamax reduced serum media (ThermoFisher Scientific) containing 10 ng/mL Polybrene Infection/Transfection Reagent (gifted from Dr Hurlstone). Colonies were selected and protein expression was then confirmed by Western blot (see Supplementary Fig. 2a). These cell lines are referred to as HeLa_FGFR2b^ST^, HeLa_FGFR2b-APEX2^ST^ and T47D_FGFR2b^KO^_FGFR2b-APEX2^ST^.

## QUANTITATIVE PHOSPHOPROTEOMICS

### Sample Preparation

For all experiments, each treatment was analysed in biological triplicates.

#### HeLa samples for phosphoproteomics

Cells were washed with PBS and lysed in ice-cold 1% triton lysis buffer supplemented with Pierce protease inhibitor tablet (Life Technologies) and phosphatase inhibitors: 5 nM Na3VO4, 5 mM NaF and 5 mM b-glycerophosphate. 5 mg of protein was obtained for each experimental condition. Proteins were precipitated overnight at -20°C in four-fold excess of ice-cold acetone. The acetone-precipitated proteins were solubilized in denaturation buffer (6 M urea, 2 M thiourea in 10 mM HEPES pH 8). Cysteines were reduced with 1 mM dithiothreitol (DTT) and alkylated with 5.5 mM chloroacetamide (CAA). Proteins were digested with endoproteinase Lys-C (Wako, Osaka, Japan) and sequencing grade modified trypsin (modified sequencing grade, Sigma) followed by quenching with 1% trifluoroacetic acid (TFA). Peptides were purified using reversed-phase Sep-Pak C18 cartridges (Waters, Milford, MA) and eluted with 50% ACN and enriched for phosphoserine-, phosphothreonine- and phosphotyrosine-containing peptides, with Titansphere chromatography. Six mL of 12% TFA ACN was added to the eluted peptides and subsequently enriched with TiO_2_ beads (5 μm, GL Sciences Inc., Tokyo, Japan). The beads were suspended in 20 mg/mL 2,5-dihydroxybenzoic acid (DHB), 80% ACN, and 6% TFA and the samples were incubated in a sample to bead ratio of 1:2 (w/w) in batch mode for 15 min with rotation. After 5 min centrifugation the beads were washed with 0% ACN, 6% TFA followeIy 40% ACN, 6% TFA and collected on C8 STAGE-tips and finally waId by 80% ACN, 6% TFA. Elution of phosphorylated peptides was done with 20ul 5% NH3 followed by 20 μL 10% NH3 in 25% ACN, which were evaporated to a final volume of 5 μL in a sped vacuum. The concentrated phosphorylated peptides were acidified with addition of 20 μI 0.1% TFA, 5% ACN and loaded on C18 STAGE-tips. Peptides were eluted from STAGE-tips in 20 μL of 40% ACN fIowIby 10 μL 60% ACN and ACN and reduced to 5 μL by SpeIac and 5 μL 0.1% FA, 5% ACN was added^22^.

#### HeLa samples for proteomics

30 ug of protein was collected as a proteome sample for in-gel digestion ^12^. Samples were prepared in lysis buffer as above containing 10 mM DTT, alkylated with 5.5 mM CAA and run on 1.00 mm Invitrogen NuPAGE 4-12% Bis-Tris Gel (Invitrogen) in NuPAGE MOPs running buffer (Invitrogen). Gels were washed in ddH_2_O, and proteins subsequently fixed and stained with Colloidal Coomassie Stain (Invitrogen). Each sample was equally separated into four fractions, and de-stained using 50% 20 mM ammonium bicarconate (NH_4_HCO_3_) + 50% ethanol (EtOH). Peptides were digested by trypsin in 50 mM NH_4_HCO_3_, neutralised and extracted using increasing concentrations of ACN, starting with 50% ACN, 3% TFA and ending with 100% ACN. Digested peptides were evaporated to a final volume of 100 μL in a speed vacuum and loaded on C18 STAGE-tips. Peptides were eluted from STAGE-tips with 20 μL of 40% ACN followed by 10 μL 60% ACN and ACN and reduced to 5 μL by SpeedVac and 5 μL 0.1% FA, 5% ACN added.

#### T47D samples for proteomics and phosphoproteomics

Cells were washed with PBS and prepared as described above up to elution off Sep-Pak C18 cartridges (Waters, Milford, MA) with 50% ACN. Prior to enrichment for phosphoserine-, phosphothreonine- and phosphotyrosine-containing peptides, with Titansphere chromatography, a small amount of the eluted peptides (1%) was taken for proteome analysis: after evaporation in a speed vacuum, peptides were resuspended in 40 μl of 0.1% TFA, 5% ACN and loaded on C18 STAGE-tips. Six mL of 12% TFA in ACN was added to the eluted peptides and subsequently enriched with TiO_2_ beads (5 μm, GL Sciences Inc., Tokyo, Japan). The beads were suspended in 20 mg/mL 2,5-dihydroxybenzoic acid (DHB), 80% ACN, and 6% TFA and the samples were incubated in a sample to bead ratio of 1:2 (w/w) in batch mode for 15 min with rotation. After 5 min centrifugation the supernatant were collected and incubated a second time with a two-fold dilution of the previous bead suspension. Sample preparation continued as described above.

#### HeLa sample for global and proximal proteomics and phosphoproteomics

Cells were pre-incubated for 40 min with Biotin Phenol (Iris Biotech) and either left untreated or treated with FGF10 (100 ng/mL) for 40 min. Hydrogen peroxide (Sigma Aldrich) was added for 1 min before quenching with Trolox (Sigma Aldrich) and sodium ascorbate (VWR) during ice cold lysis. Cells were lysed using APEX-RIPA buffer (50mM Tris-HCl pH 7.5. 150mM NaCl. 0.1% SDS. 1% Triton. 0.5% sodium deoxycholate. 1x protease inhibitor cocktail. 1mM PMSF. 10mM sodium ascorbate. 10mM sodium azide. 5mM Trolox) and protein extracted as described above. For each sample, 15 mg of protein was collected. Of this amount,120 μg was taken and run on a gradient gel to acquire the global proteome; 5 mg was precipitated in acetone and processed as described above to obtain the global phosphoproteome; the rest of the lysate was enriched for biotinylation using a 2 h room temperature pull-down with streptavidin beads, to generate the proximal proteome. A fifth of the bead slurry was stripped using boiling 4x sample buffer enrich with biotin, the supernatant was run on a gradient gel to acquire the proximal proteome. The remaining streptavidin bead slurry was stripped using boiling 8M guanidine ph1.5 supplemented in 5mM TCEP and 10mM CAA. Reduced samples where then difested with LysC for 60min RT, diluted to 1M guanidine using Tris 25mM pH8.5, before digestion with trypsin and enrichment for phosphorylated peptides as described above.

### Mass Spectrometry

Purified peptides were analysed by LC-MS/MS using an UltiMate® 3000 Rapid Separation LC (RSLC, Dionex Corporation, Sunnyvale, CA) coupled to a QE-HF (Thermo Fisher Scientific, Waltham, MA) mass spectrometer^22^. Mobile phase A was 0.1% FA in water and mobile phase B was 0.1% FA in ACN and the column was a 75 mm x 250 μm inner diameter 1.7 mM CSH C18, analytical column (Waters). A 1μl aliquot of the sample (for proteome analysis) or a 3μl aliquot was transferred to a 5μl loop and loaded on to the column at a flow of 300nl/min at 5% B for 5 and 13 min, respectively. The loop was then taken out of line and the flow was reduced from 300nl/min to 200nl/min in 1 min, and to 7% B. Peptides were separated using a gradient that went from 7% to 18% B in 64 min, then from 18% to 27% B in 8 min and finally from 27% B to 60% B in 1 min. The column was washed at 60% B for 3 min and then REs-equilibrated for a further 6.5 min. At 85 min the flow was increased to 300nl/min until the end of the run at 90min. Mass spectrometry data was acquired in a data directed manner for 90 min in positive mode. Peptides were selected for fragmentation automatically by data dependent analysis on a basis of the top 8 (phosphoproteome analysis) or top 12 (proteome analysis) with m/z between 300 to 1750Th and a charge state of 2, 3 or 4 with a dynamic exclusion set at 15 sec. The MS Resolution was set at 120,000 with an AGC target of 3e6 and a maximum fill time set at 20ms. The MS2 Resolution was set to 60,000, with an AGC target of 2e5, and a maximum fill time of 110 ms for Top12 methods, and 30,000, with an AGC target of 2e5, and a maximum fill time of 45 ms for Top8 analysis. The isolation window was of 1.3Th and the collision energy was of 28.

### Raw Files Analysis

Raw data were analysed by the MaxQuant software suite (https://www.maxquant.org; version 1.6.2.6 and 1.5.6.5) using the integrated Andromeda search engine^85^. Proteins were identified by searching the HCD-MS/MS peak lists against a target/decoy version of the human Uniprot Knowledgebase database that consisted of the complete proteome sets and isoforms (v.2019; https.//uniprot.org/proteomes/UP000005640_9606) supplemented with commonly observed contaminants such as porcine trypsin and bovine serum proteins. Tandem mass spectra were initially matched with a mass tolerance of 7 ppm on precursor masses and 0.02 Da or 20 ppm for fragment ions. Cysteine carbamidomethylation was searched as a fixed modification. Protein N-acetylation, N-pyro-glutamine, oxidized methionine, and phosphorylation of serine, threonine, and tyrosine were searched as variable modifications for the phosphoproteomes. Protein N-acetylation, oxidized methionine and deamidation of asparagine and glutamine were searched as variable modifications for the proteome experiments. Biotinylation by BP (Y; (C18H23N3O3S)) was included as a variable and fixed modification for <u>global and proximal</u> raw files. Label-free parameters were used for all the analysis as described^86^. False discovery rate was set to 0.01 for peptides, proteins, and modification sites. Minimal peptide length was six amino acids. Site localization probabilities were calculated by MaxQuant using the PTM scoring algorithm^39^.The dataset were filtered by posterior error probability to achieve a false discovery rate below 1% for peptides, proteins and modification sites. Only peptides with Andromeda score >40 were included.

### Data and Statistical Analysis

All statistical and bioinformatics analyses were done using the freely available software Perseus, version 1.6.5.0 or 1.6.2.1.^87^, R framework and Bioconductor^88^, Python framework (available at http://www.python.org), SubcellulaRVis (https://www.biorxiv.org/content/10.1101/2021.11.18.469118v1), STRING v11.5^89^, Cytoscape (version 3.7.2)^90^. Over-representation analysis (ORA) of KEGG terms was performed using Enrichr and the EnrichR R inferface^91^.

All measured peptide intensities were normalized using the “normalizeQuantiles” function from the Bioconductor R-package *limma*. Potential contaminant proteins or phosphorylated-peptides and peptides or phosphorylated-peptides matching the reverse sequence database were removed^22^. For all datasets, phosphorylated peptides with a localization score greater than 0.75 were included in the downstream bioinformatics analysis. Pearson correlation was calculated in R.

For both the HeLa and the T47D phosphoproteomics datasets, samples were normalized separately (Smith et al., 2021) and were grouped based on treatment and only phosphorylated peptides with values in all three replicates of at least one treatment were included in further analysis. Missing values were subsequently imputed from a normal distribution using Perseus default settings. Median z-score of intensities were used for further analysis. The HeLa-FGFR2b dataset was separated into eleven clusters by fuzzy c-means clustering using the “fanny” function from the R package “cluster” performed after multi-sample ANOVA test with FDR > 0.0001 in Perseus. The clustering results of the HeLa-FGFR2b dataset were then used as a training dataset for the classification of phosphorylated sites of the T47D cell line. Kernelized Parzen window (i.e. kernel density estimation) classifier scripted via Python library “statsmodels” (version: 0.11.1) was used as a supervised learning method for generating the classification results. Over-representation analysis (ORA) of KEGG terms was performed using Enrichr and the EnrichR R inferface^91^ and significantly over-represented terms within the data were represented in dot plots. The SubCellulaRVis tool (https://www.biorxiv.org/content/10.1101/2021.11.18.469118v1) was used to visualize the sub-cellular localization of proteins.

For the SRP datasets, phosphorylated site intensities acquired from global and proximal MS runs were normalized separately. Replicates were summarised by calculating the median of normalized intensities. Missing values were imputed using random draws from a truncated distribution with the *impute.QRLIC()* function from the R CRAN package *imputeLCMD.* PCA, Students’ t-test and one-way ANOVA were calculated using the prcomp(), t.test() and aov() functions in R, respectively. Hierarchical clustering was performed using the *hclust()* function in R. ORA and use of SubCellulaRVis tool were performed as described above. Networks were constructed using STRING; only interactions with an experimental confidence > 0.4 were included. KEGG pathways were extracted using the *EnrichmentBrowser* package from Bioconductor.

## BIOCHEMICAL AND FUNCTIONAL ASSAYS

### Cell Lysis and Western Blotting

Cells were serum starved overnight in serum-free medium and stimulated for the indicated time points with 100 ng/mL of FGF7 or FGF10^13^. Where indicated, cells were pre-incubated for 2 h with 100 nM PD173074 (Selleckchem, #S1264), 0.5 μM Rapamycin (Sigma Aldrich, #37094), 2 μM ULK101 (Selleckchem, #S8793), 10 μM SBI-0206965 (Selleckchem, #S7885), 10 μM Dynasore (Abcam, #ab120192), Monensin sodium salt, Na^+^ ionophore (Abcam, #ab120499) and 200 μM Primaquine bisphosphate (Sigma Aldrich, #160393). Control cells were pre-incubated with DMSO alone. Cells treated with glucose-removal, Glucose-6-phosphate (Sigma-Aldrich, #10127647001), Sodium valporate (Sigma-Aldrich, #BP452) and Fluspiriline (Sigma-Aldrich, #F100) were treated for 24 h prior to FGF10 stimulation. After stimulation, cell extraction and immunoblotting were performed as previously described ^22^. Proteins were resolved by SDS-PAGE and transferred to nitrocellulose membranes (Protran, Biosciences). Proteins of interest were visualized using specific antibodies, followed by peroxidase-conjugated secondary antibodies and by an enhanced chemiluminescence kit (Amersham Biosciences). Blots were visualised using the Universal Hood II Gel Molecular Imaging System (Bio-Rad). Each experiment was repeated at least three times and produced similar results.

### Biotinylation assays

Biotinylation pull downs experiments were performed as described previously^22^. Briefly, cells were pre-incubated (40 min) with Biotin Phenol (Iris Biotech) after stimulation with ligands. Hydrogen peroxide (Sigma Aldrich) was added for 1 min before quenching with Trolox (Sigma Aldrich) and Sodium ascorbate (VWR) during ice cold lysis. A 2 h RT pull-down with streptavidin beads was then performed and the supernatant was run against the bound proteins and the total lysates.

### RNA Isolation and real-time qPCR analysis

RNA from cell lines was isolated with TRIZOL® (Invitrogen). After chloroform extraction and centrifugation, 5 µg RNA was DNase treated using RNase-Free DNase Set (Qiagen) and 1 µg of DNase treated RNA was then taken for cDNA synthesis using the Protoscript I first strand cDNA synthesis kit (New England Biolabs). Selected genes were amplified by quantitative real time PCR (RT-qPCR) using Sygreen (PCR Biosystems). Relative expression was calculated using the delta-delta CT methodology and beta-actin was used as reference housekeeping gene. Sequences for primers used can be found in the key resource table. qPCR machine used was Applied Biosystems MX300P.

**Table 3:**
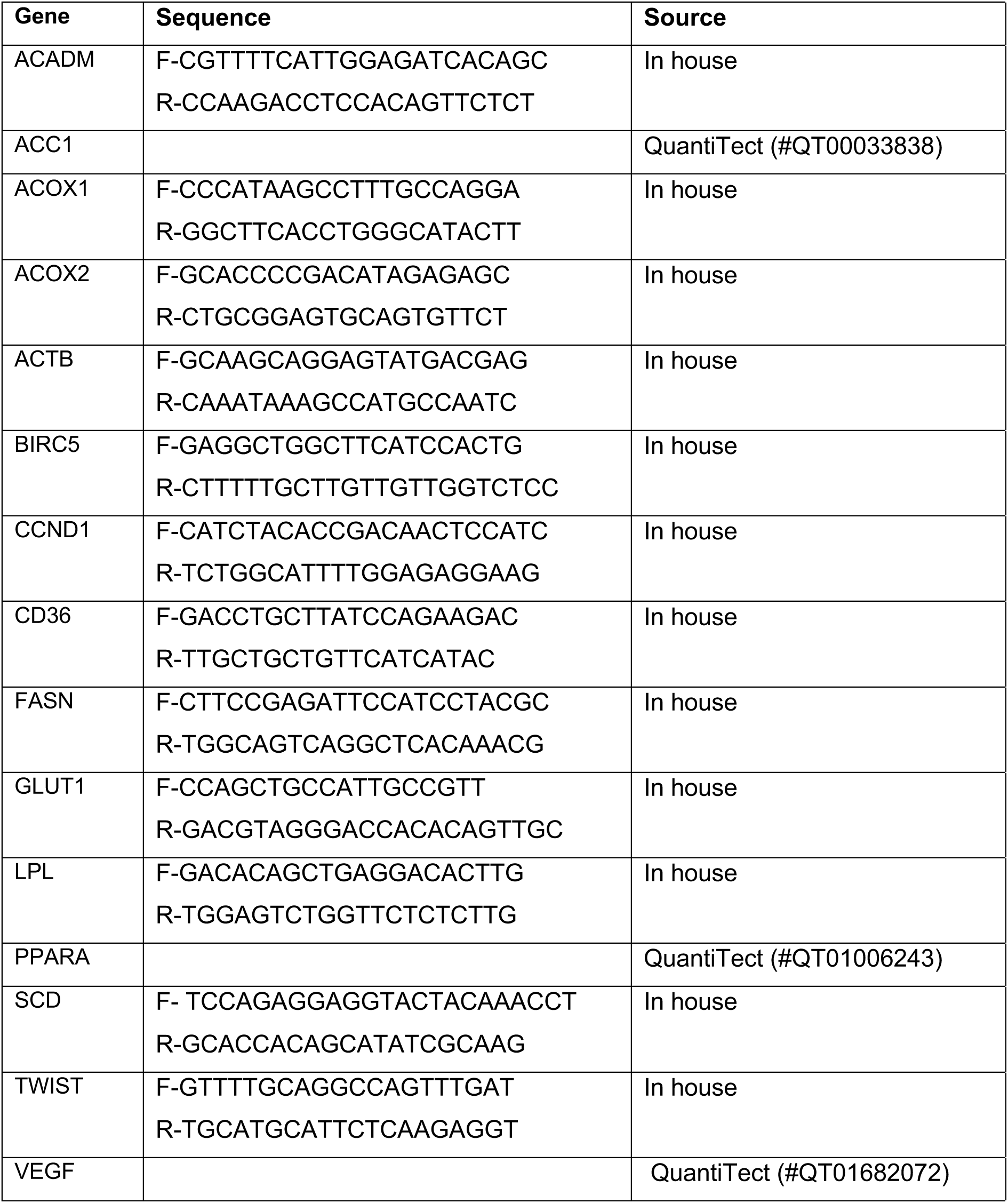
Primers for RT-qPCR

### Proliferation Assays

#### EdU Incorporation

Indicated cells were labelled with 20 µM 5-ethynyl-2’-deoxyuridine (EdU) for 4 h and processed following the manufacturer’s protocol (Click-iT® EdU Alexa Fluor® 488 Imaging Kit, Thermo Fisher). Prior to imaging cells were then stained with 5ng/ml Hoecsht 3342 for 15 min. Stained cells were analysed using a using a Leica microscope system. Statistical analysis was performed at the endpoint across repeats, as indicated in the Figure legends.

#### Cleaved caspase assay

Apoptosis was measured in cells receiving either 24 h treatment with FGF10. Appropriately treated cells were incubated with 20 mM CellEvent™ Caspase-3/7 Green Detection Reagent (Invitrogen) made to 100X in PBS for 4 h in darkness then washed thoroughly in 1X PBS. Fluorescence was measured at 502 nm excitation and 530 nm emission. Statistical analysis was performed at the endpoint across repeats, as indicated in the Figure legends.

#### Autophagy

Cells were assayed for autophagy using 5 mM Acridine Orange (Sigma) for 30 min after which excess was removed by thorough washing with 1X PBS. This fluorophore appears green when diffuse but is shifted to the red end of the spectrum when accumulated in acidic vesicles ^92^. As such, excitation/emission wavelengths of 500/526 nm were used to measure intensity of diffuse acridine orange (non-specific) and 460/650 nm to assess autophagic staining. The ratio of these values represents stained autophagosomes. Statistical analysis was performed at the endpoint across repeats, as indicated in the Figure legends.

## IMMUNOFLUORESCENCE and QUANTIFICATION

Immunofluorescence staining was performed as previously described ^22^. To detect FGFR2 cells were incubated with anti-FGFR2 antibody (Cell Signalling) for 45 min with gentle agitation. The binding of the antibody did not activate receptor signalling in untreated cells nor induced receptor internalization as previously reported ^22^. After stimulation cells were incubated at 37°C for different time points. At each time point, non-permeabilized cells were either fixed to visualize the receptor on the cell surface (plasma membrane) or acid-washed in ice-cold buffer (50 mM glycine, pH 2.5) to remove surface-bound antibody. Acid-washed cells were then fixed and permeabilized to visualize the internalized receptor (cytoplasm). Finally, to detect FGFR2b cells were stained with AlexaFluor488-conjugated donkey anti-mouse or anti-rabbit (Jackson ImmunoResearch Laboratories). Nuclei were stained with DAPI. Coverslips were then mounted in mounting medium (Vectashield; Vector Laboratories).

For co-localization experiments, cells were acid washed, fixed, permeabilized with 0.02% saponin (Sigma), treated with the indicated primary antibody for 60 min at 37 °C, and stained with AlexaFluor488 (or 568 or 647)-conjugated donkey anti-mouse or anti-rabbit. Samples expressing GFP-tagged proteins were kept in the dark. Nuclei were stained with DAPI. Coverslips were then mounted in mounting medium (Vectashield; Vector Laboratories).

All the images were acquired at room temperature on a Leica TCS SP8 AOBS inverted confocal using a 100x oil immersion objective and 2.5x or 3x confocal zoom. The confocal settings were as follows: pinhole, 1 airy unit, format, 1024 x 1024. Images were collected using the following detection mirror settings: FITC 494-530nm; Texas red 602-665nm; Cy5 640-690nm. The images were collected sequentially. Raw images were exported as .lsm files, and adjustments in image contrast and brightness were applied identical for all images in a given experiment using the freely available software Image J v. 1.52p^93^.

Quantification of FGFR2b recycling, Co-localization, and Expression Fraction was performed as recently described in detail^22^. The scripts for the quantification of co-localization were written in the Python language and the code for Costes-adjusted MCC was taken verbatim from the CellProfiler code base. Statistical analysis was performed across repeats, as indicated in the Figure legends.

## DATA AVAILABILITY

## REFERENCES

1. Cullen, P. J. & Steinberg, F. To degrade or not to degrade: mechanisms and significance of endocytic recycling. Nat Rev Mol Cell Biol 19, 679–696, doi:10.1038/s41580-018-0053-7 (2018).

2. Goh, L. K. & Sorkin, A. Endocytosis of receptor tyrosine kinases. Cold Spring Harb Perspect Biol 5, a017459, doi:10.1101/cshperspect.a017459 (2013).

3. MacDonald, E., Savage, B. & Zech, T. Connecting the dots: combined control of endocytic recycling and degradation. Biochem Soc Trans 48, 2377–2386, doi:10.1042/BST20180255 (2020).

4. Hsu, V. W., Bai, M. & Li, J. Getting active: protein sorting in endocytic recycling. Nat Rev Mol Cell Biol 13, 323–328, doi:10.1038/nrm3332 (2012).

5. Sigismund, S., Lanzetti, L., Scita, G. & Di Fiore, P. P. Endocytosis in the context-dependent regulation of individual and collective cell properties. Nat Rev Mol Cell Biol 22, 625–643, doi:10.1038/s41580-021-00375-5 (2021).

6. Miaczynska, M. Effects of membrane trafficking on signaling by receptor tyrosine kinases. Cold Spring Harb Perspect Biol 5, a009035, doi:10.1101/cshperspect.a009035 (2013).

7. Naslavsky, N. & Caplan, S. The enigmatic endosome - sorting the ins and outs of endocytic trafficking. J Cell Sci 131, doi:10.1242/jcs.216499 (2018).

8. O’Sullivan, M. J. & Lindsay, A. J. The Endosomal Recycling Pathway-At the Crossroads of the Cell. Int J Mol Sci 21, doi:10.3390/ijms21176074 (2020).

9. Schmid, S. L. Reciprocal regulation of signaling and endocytosis: Implications for the evolving cancer cell. J Cell Biol 216, 2623–2632, doi:10.1083/jcb.201705017 (2017).

10. Stasyk, T. & Huber, L. A. Spatio-Temporal Parameters of Endosomal Signaling in Cancer: Implications for New Treatment Options. J Cell Biochem 117, 836–843, doi:10.1002/jcb.25418 (2016).

11. Caswell, P. T. et al. Rab-coupling protein coordinates recycling of alpha5beta1 integrin and EGFR1 to promote cell migration in 3D microenvironments. J Cell Biol 183, 143–155, doi:10.1083/jcb.200804140 (2008).

12. Francavilla, C. et al. Multilayered proteomics reveals molecular switches dictating ligand-dependent EGFR trafficking. Nat Struct Mol Biol 23, 608–618, doi:10.1038/nsmb.3218 (2016).

13. Francavilla, C. et al. Functional proteomics defines the molecular switch underlying FGF receptor trafficking and cellular outputs. Mol Cell 51, 707–722, doi:10.1016/j.molcel.2013.08.002 (2013).

14. Lanzetti, L. & Di Fiore, P. P. Behind the Scenes: Endo/Exocytosis in the Acquisition of Metastatic Traits. Cancer Res 77, 1813–1817, doi:10.1158/0008-5472.CAN-16-3403 (2017).

15. Yarwood, R., Hellicar, J., Woodman, P. G. & Lowe, M. Membrane trafficking in health and disease. Dis Model Mech 13, doi:10.1242/dmm.043448 (2020).

16. Wang, Y., Pennock, S., Chen, X. & Wang, Z. Endosomal signaling of epidermal growth factor receptor stimulates signal transduction pathways leading to cell survival. Mol Cell Biol 22, 7279–7290, doi:10.1128/MCB.22.20.7279-7290.2002 (2002).

17. Teis, D., Wunderlich, W. & Huber, L. A. Localization of the MP1-MAPK scaffold complex to endosomes is mediated by p14 and required for signal transduction. Dev Cell 3, 803–814, doi:10.1016/s1534-5807(02)00364-7 (2002).

18. Bruggemann, Y., Karajannis, L. S., Stanoev, A., Stallaert, W. & Bastiaens, P. I. H. Growth factor-dependent ErbB vesicular dynamics couple receptor signaling to spatially and functionally distinct Erk pools. Sci Signal 14, doi:10.1126/scisignal.abd9943 (2021).

19. Saxton, R. A. & Sabatini, D. M. mTOR Signaling in Growth, Metabolism, and Disease. Cell 168, 960–976, doi:10.1016/j.cell.2017.02.004 (2017).

20. Savini, M., Zhao, Q. & Wang, M. C. Lysosomes: Signaling Hubs for Metabolic Sensing and Longevity. Trends Cell Biol 29, 876–887, doi:10.1016/j.tcb.2019.08.008 (2019).

21. Yuan, W. & Song, C. The Emerging Role of Rab5 in Membrane Receptor Trafficking and Signaling Pathways. Biochem Res Int 2020, 4186308, doi:10.1155/2020/4186308 (2020).

22. Smith, M. P. et al. Reciprocal priming between receptor tyrosine kinases at recycling endosomes orchestrates cellular signalling outputs. EMBO J 40, e107182, doi:10.15252/embj.2020107182 (2021).

23. Szybowska, P., Kostas, M., Wesche, J., Haugsten, E. M. & Wiedlocha, A. Negative Regulation of FGFR (Fibroblast Growth Factor Receptor) Signaling. Cells 10, doi:10.3390/cells10061342 (2021).

24. Ornitz, D. M. & Itoh, N. The Fibroblast Growth Factor signaling pathway. Wiley Interdiscip Rev Dev Biol 4, 215–266, doi:10.1002/wdev.176 (2015).

25. Ferguson, H. R., Smith, M. P. & Francavilla, C. Fibroblast Growth Factor Receptors (FGFRs) and Noncanonical Partners in Cancer Signaling. Cells 10, doi:10.3390/cells10051201 (2021).

26. Watson, J. & Francavilla, C. Regulation of FGF10 Signaling in Development and Disease. Front Genet 9, 500, doi:10.3389/fgene.2018.00500 (2018).

27. Belleudi, F. et al. Keratinocyte growth factor receptor ligands target the receptor to different intracellular pathways. Traffic 8, 1854–1872, doi:10.1111/j.1600-0854.2007.00651.x (2007).

28. Lobingier, B. T. et al. An Approach to Spatiotemporally Resolve Protein Interaction Networks in Living Cells. Cell 169, 350–360 e312, doi:10.1016/j.cell.2017.03.022 (2017).

29. Paek, J. et al. Multidimensional Tracking of GPCR Signaling via Peroxidase-Catalyzed Proximity Labeling. Cell 169, 338–349 e311, doi:10.1016/j.cell.2017.03.028 (2017).

30. Markmiller, S. et al. Context-Dependent and Disease-Specific Diversity in Protein Interactions within Stress Granules. Cell 172, 590–604 e513, doi:10.1016/j.cell.2017.12.032 (2018).

31. Han, S. et al. Proximity Biotinylation as a Method for Mapping Proteins Associated with mtDNA in Living Cells. Cell Chem Biol 24, 404–414, doi:10.1016/j.chembiol.2017.02.002 (2017).

32. Han, Y. et al. Directed Evolution of Split APEX2 Peroxidase. ACS Chem Biol 14, 619–635, doi:10.1021/acschembio.8b00919 (2019).

33. Gingras, A. C., Abe, K. T. & Raught, B. Getting to know the neighborhood: using proximity-dependent biotinylation to characterize protein complexes and map organelles. Curr Opin Chem Biol 48, 44–54, doi:10.1016/j.cbpa.2018.10.017 (2019).

34. Go, C. D. et al. A proximity-dependent biotinylation map of a human cell. Nature 595, 120–124, doi:10.1038/s41586-021-03592-2 (2021).

35. Zellner, S., Schifferer, M. & Behrends, C. Systematically defining selective autophagy receptor-specific cargo using autophagosome content profiling. Mol Cell 81, 1337–1354 e1338, doi:10.1016/j.molcel.2021.01.009 (2021).

36. Francavilla, C. et al. The binding of NCAM to FGFR1 induces a specific cellular response mediated by receptor trafficking. J Cell Biol 187, 1101–1116, doi:10.1083/jcb.200903030 (2009).

37. Surve, S., Watkins, S. C. & Sorkin, A. EGFR-RAS-MAPK signaling is confined to the plasma membrane and associated endorecycling protrusions. J Cell Biol 220, doi:10.1083/jcb.202107103 (2021).

38. Schindler, C., Chen, Y., Pu, J., Guo, X. & Bonifacino, J. S. EARP is a multisubunit tethering complex involved in endocytic recycling. Nat Cell Biol 17, 639–650, doi:10.1038/ncb3129 (2015).

39. Olsen, J. V. et al. Global, in vivo, and site-specific phosphorylation dynamics in signaling networks. Cell 127, 635–648, doi:10.1016/j.cell.2006.09.026 (2006).

40. Shin, J. H., Park, C. W., Yoon, G., Hong, S. M. & Choi, K. Y. NNMT depletion contributes to liver cancer cell survival by enhancing autophagy under nutrient starvation. Oncogenesis 7, 58, doi:10.1038/s41389-018-0064-4 (2018).

41. Klionsky, D. J. et al. Guidelines for the use and interpretation of assays for monitoring autophagy (4th edition)(1). Autophagy 17, 1–382, doi:10.1080/15548627.2020.1797280 (2021).

42. Ichimiya, T. et al. Autophagy and Autophagy-Related Diseases: A Review. Int J Mol Sci 21, doi:10.3390/ijms21238974 (2020).

43. Schaaf, M. B., Keulers, T. G., Vooijs, M. A. & Rouschop, K. M. LC3/GABARAP family proteins: autophagy-(un)related functions. FASEB J 30, 3961–3978, doi:10.1096/fj.201600698R (2016).

44. Ashkenazi, A. et al. Polyglutamine tracts regulate beclin 1-dependent autophagy. Nature 545, 108–111, doi:10.1038/nature22078 (2017).

45. Xia, H. G. et al. Control of basal autophagy by calpain1 mediated cleavage of ATG5. Autophagy 6, 61–66, doi:10.4161/auto.6.1.10326 (2010).

46. Xia, Q. et al. Valproic acid induces autophagy by suppressing the Akt/mTOR pathway in human prostate cancer cells. Oncol Lett 12, 1826–1832, doi:10.3892/ol.2016.4880 (2016).

47. Resat, H., Ewald, J. A., Dixon, D. A. & Wiley, H. S. An integrated model of epidermal growth factor receptor trafficking and signal transduction. Biophys J 85, 730–743, doi:10.1016/s0006-3495(03)74516-0 (2003).

48. Zachari, M. & Ganley, I. G. The mammalian ULK1 complex and autophagy initiation. Essays Biochem 61, 585–596, doi:10.1042/EBC20170021 (2017).

49. Longatti, A. & Tooze, S. A. Recycling endosomes contribute to autophagosome formation. Autophagy 8, 1682–1683, doi:10.4161/auto.21486 (2012).

50. Macia, E. et al. Dynasore, a cell-permeable inhibitor of dynamin. Dev Cell 10, 839–850, doi:10.1016/j.devcel.2006.04.002 (2006).

51. van Weert, A. W., Geuze, H. J., Groothuis, B. & Stoorvogel, W. Primaquine interferes with membrane recycling from endosomes to the plasma membrane through a direct interaction with endosomes which does not involve neutralisation of endosomal pH nor osmotic swelling of endosomes. Eur J Cell Biol 79, 394–399, doi:10.1078/0171-9335-00062 (2000).

52. Kim, Y. M. et al. SH3BP4 is a negative regulator of amino acid-Rag GTPase-mTORC1 signaling. Mol Cell 46, 833–846, doi:10.1016/j.molcel.2012.04.007 (2012).

53. Budayeva, H. G. & Kirkpatrick, D. S. Monitoring protein communities and their responses to therapeutics. Nat Rev Drug Discov 19, 414–426, doi:10.1038/s41573-020-0063-y (2020).

54. Wang, W., Bian, J. & Li, Z. Internalized Activation of Membrane Receptors: From Phenomenon to Theory. Trends Cell Biol 31, 428–431, doi:10.1016/j.tcb.2021.03.008 (2021).

55. Barrow-McGee, R. & Kermorgant, S. Met endosomal signalling: in the right place, at the right time. Int J Biochem Cell Biol 49, 69–74, doi:10.1016/j.biocel.2014.01.009 (2014).

56. Lopez-Otin, C. & Kroemer, G. Hallmarks of Health. Cell 184, 33–63, doi:10.1016/j.cell.2020.11.034 (2021).

57. Chernikova, S. B. et al. Dynamin impacts homology-directed repair and breast cancer response to chemotherapy. J Clin Invest 128, 5307–5321, doi:10.1172/JCI87191 (2018).

58. Liu, Y. et al. Spatiotemporally resolved subcellular phosphoproteomics. Proc Natl Acad Sci U S A 118, doi:10.1073/pnas.2025299118 (2021).

59. Qin, W., Myers, S. A., Carey, D. K., Carr, S. A. & Ting, A. Y. Spatiotemporally-resolved mapping of RNA binding proteins via functional proximity labeling reveals a mitochondrial mRNA anchor promoting stress recovery. Nat Commun 12, 4980, doi:10.1038/s41467-021-25259-2 (2021).

60. Bosch, J. A., Chen, C. L. & Perrimon, N. Proximity-dependent labeling methods for proteomic profiling in living cells: An update. Wiley Interdiscip Rev Dev Biol 10, e392, doi:10.1002/wdev.392 (2021).

61. Cristea, I. M. & Lilley, K. S. Editorial overview: Untangling proteome organization in space and time. Curr Opin Chem Biol 48, A1–A4, doi:10.1016/j.cbpa.2019.02.001 (2019).

62. Lundberg, E. & Borner, G. H. H. Spatial proteomics: a powerful discovery tool for cell biology. Nat Rev Mol Cell Biol 20, 285–302, doi:10.1038/s41580-018-0094-y (2019).

63. Christopher, J. A. et al. Subcellular proteomics. Nat Rev Methods Primers 1, doi:10.1038/s43586-021-00029-y (2021).

64. Thapa, N. et al. Phosphatidylinositol-3-OH kinase signalling is spatially organized at endosomal compartments by microtubule-associated protein 4. Nat Cell Biol 22, 1357–1370, doi:10.1038/s41556-020-00596-4 (2020).

65. Jiang, T., Pan, C. Q. & Low, B. C. BPGAP1 spatially integrates JNK/ERK signaling crosstalk in oncogenesis. Oncogene 36, 3178–3192, doi:10.1038/onc.2016.466 (2017).

66. Dauner, K., Eid, W., Raghupathy, R., Presley, J. F. & Zha, X. mTOR complex 1 activity is required to maintain the canonical endocytic recycling pathway against lysosomal delivery. J Biol Chem 292, 5737–5747, doi:10.1074/jbc.M116.771451 (2017).

67. Kvainickas, A. et al. Retromer and TBC1D5 maintain late endosomal RAB7 domains to enable amino acid-induced mTORC1 signaling. J Cell Biol 218, 3019–3038, doi:10.1083/jcb.201812110 (2019).

68. Takahashi, Y. et al. The late endosome/lysosome-anchored p18-mTORC1 pathway controls terminal maturation of lysosomes. Biochem Biophys Res Commun 417, 1151–1157, doi:10.1016/j.bbrc.2011.12.082 (2012).

69. Miller, K., Beardmore, J., Kanety, H., Schlessinger, J. & Hopkins, C. R. Localization of the epidermal growth factor (EGF) receptor within the endosome of EGF-stimulated epidermoid carcinoma (A431) cells. J Cell Biol 102, 500–509, doi:10.1083/jcb.102.2.500 (1986).

70. Fraser, J., Cabodevilla, A. G., Simpson, J. & Gammoh, N. Interplay of autophagy, receptor tyrosine kinase signalling and endocytic trafficking. Essays Biochem 61, 597–607, doi:10.1042/EBC20170091 (2017).

71. Fraser, J. et al. Targeting of early endosomes by autophagy facilitates EGFR recycling and signalling. EMBO Rep 20, e47734, doi:10.15252/embr.201947734 (2019).

72. Nanni, M., Ranieri, D., Rosato, B., Torrisi, M. R. & Belleudi, F. Role of Fibroblast Growth Factor Receptor 2b in the Cross Talk between Autophagy and Differentiation: Involvement of Jun N-Terminal Protein Kinase Signaling. Mol Cell Biol 38, doi:10.1128/MCB.00119-18 (2018).

73. Cinque, L. et al. FGF signalling regulates bone growth through autophagy. Nature 528, 272–275, doi:10.1038/nature16063 (2015).

74. Wu, M. & Zhang, P. EGFR-mediated autophagy in tumourigenesis and therapeutic resistance. Cancer Lett 469, 207–216, doi:10.1016/j.canlet.2019.10.030 (2020).

75. Birgisdottir, A. B. & Johansen, T. Autophagy and endocytosis - interconnections and interdependencies. J Cell Sci 133, doi:10.1242/jcs.228114 (2020).

76. Wang, X. et al. Endocytosis and Organelle Targeting of Nanomedicines in Cancer Therapy. Int J Nanomedicine 15, 9447–9467, doi:10.2147/IJN.S274289 (2020).

77. Yadav, V., Tolwinski, N. & Saunders, T. E. Spatiotemporal sensitivity of mesoderm specification to FGFR signalling in the Drosophila embryo. Sci Rep 11, 14091, doi:10.1038/s41598-021-93512-1 (2021).

78. Mathiassen, S. G., De Zio, D. & Cecconi, F. Autophagy and the Cell Cycle: A Complex Landscape. Front Oncol 7, 51, doi:10.3389/fonc.2017.00051 (2017).

79. Nowosad, A. et al. p27 controls Ragulator and mTOR activity in amino acid-deprived cells to regulate the autophagy-lysosomal pathway and coordinate cell cycle and cell growth. Nat Cell Biol 22, 1076–1090, doi:10.1038/s41556-020-0554-4 (2020).

80. Yamasaki, A., Jin, Y. & Ohsumi, Y. Mitotic phosphorylation of the ULK complex regulates cell cycle progression. PLoS Biol 18, e3000718, doi:10.1371/journal.pbio.3000718 (2020).

81. Tan, C. et al. Cell size homeostasis is maintained by CDK4-dependent activation of p38 MAPK. Dev Cell 56, 1756–1769 e1757, doi:10.1016/j.devcel.2021.04.030 (2021).

82. Aveic, S. et al. Autophagy inhibition improves the cytotoxic effects of receptor tyrosine kinase inhibitors. Cancer Cell Int 18, 63, doi:10.1186/s12935-018-0557-4 (2018).

83. Porebska, N. et al. Targeting Cellular Trafficking of Fibroblast Growth Factor Receptors as a Strategy for Selective Cancer Treatment. J Clin Med 8, doi:10.3390/jcm8010007 (2018).

84. Li, Y. et al. FGFR-inhibitor-mediated dismissal of SWI/SNF complexes from YAP-dependent enhancers induces adaptive therapeutic resistance. Nat Cell Biol 23, 1187–1198, doi:10.1038/s41556-021-00781-z (2021).

85. Cox, J. et al. Andromeda: a peptide search engine integrated into the MaxQuant environment. J Proteome Res 10, 1794–1805, doi:10.1021/pr101065j (2011).

86. Cox, J. et al. Accurate proteome-wide label-free quantification by delayed normalization and maximal peptide ratio extraction, termed MaxLFQ. Mol Cell Proteomics 13, 2513–2526, doi:10.1074/mcp.M113.031591 (2014).

87. Tyanova, S. & Cox, J. Perseus: A Bioinformatics Platform for Integrative Analysis of Proteomics Data in Cancer Research. Methods Mol Biol 1711, 133–148, doi:10.1007/978-1-4939-7493-1_7 (2018).

88. Huber, W. et al. Orchestrating high-throughput genomic analysis with Bioconductor. Nat Methods 12, 115–121, doi:10.1038/nmeth.3252 (2015).

89. Szklarczyk, D. et al. STRING v11: protein-protein association networks with increased coverage, supporting functional discovery in genome-wide experimental datasets. Nucleic Acids Res 47, D607–D613, doi:10.1093/nar/gky1131 (2019).

90. Shannon, P. et al. Cytoscape: a software environment for integrated models of biomolecular interaction networks. Genome Res 13, 2498–2504, doi:10.1101/gr.1239303 (2003).

91. Kuleshov, M. V. et al. Enrichr: a comprehensive gene set enrichment analysis web server 2016 update. Nucleic Acids Res 44, W90–97, doi:10.1093/nar/gkw377 (2016).

92. SenthilKumar, G., Skiba, J. H. & Kimple, R. J. High-throughput quantitative detection of basal autophagy and autophagic flux using image cytometry. Biotechniques 67, 70–73, doi:10.2144/btn-2019-0044 (2019).

93. Schindelin, J. et al. Fiji: an open-source platform for biological-image analysis. Nat Methods 9, 676–682, doi:10.1038/nmeth.2019 (2012).

