## Supplementary Information for "Spatially Resolved Phosphoproteomics Reveals Fibroblast Growth Factor Receptor Recycling-driven Regulation of Autophagy and Survival"

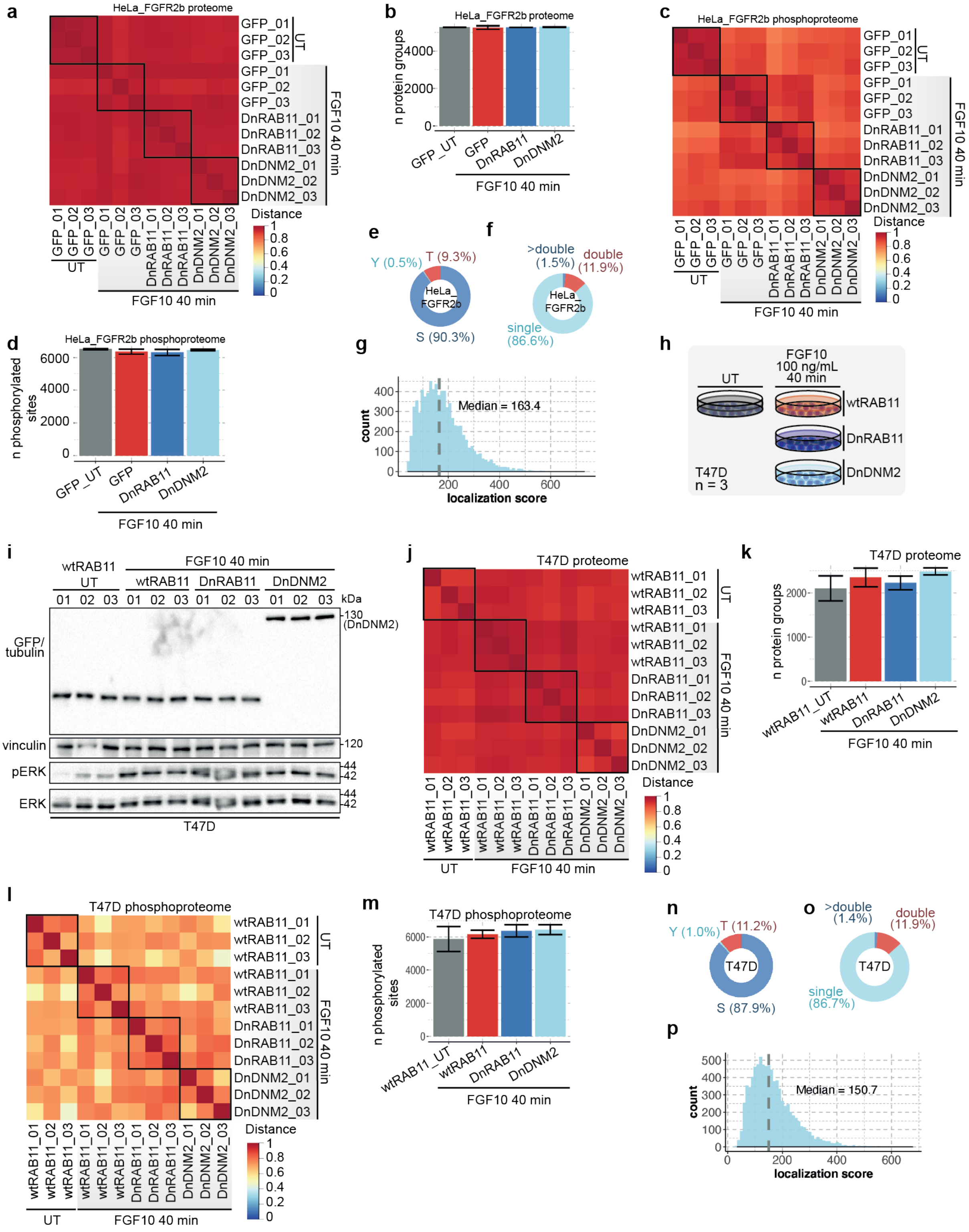


**Supplementary Fig. 1. Quality assessment of the HeLa and T47D phosphoproteome and proteome. a** Heat map showing the Pearson correlation of the HeLa proteome shows that transient expression (< 72h) of dominant negative constructs did not noticeably change the background proteome of HeLa cells. **b** Number (n) of protein groups quantified in each condition. Values represent mean ± SD of N = 3. **c** Heat map showing the Pearson correlation of the HeLa phosphoproteome shows good reproducibility and variation within experimental conditions. **d** Number (n) of phosphorylated peptides in each condition. Values represent mean ± SD of n = 3. **e** Distribution of peptides containing phosphorylated Serine (S), Threonine (T) and Tyrosine (Y) residues (%). **f** Distribution of peptides containing a single, double of more than double modification (%). **g** The distribution of phosphorylated peptides score showed that most of the peptides were identified with high Andromeda score (median: 163), consistent with [22]. **h** Workflow of the phosphoproteomics experiment in T47D cells transiently expressing either wtRAB11, DnRAB11 or DnDNM2 and left untreated (UT) or treated with FGF10 for 40 min. **i** Immunoblot analysis with the indicated antibodies of T47D cells expressing GFP and left untreated (UT) and expressing either GFP, DnRAB11 or DnDNM2 and treated with FGF10 for 40 min. **j** Heat map showing the Pearson correlation of the T47D proteome shows that transient expression (< 72h) of dominant negative constructs did not noticeably change the background proteome of T47D cells. **k** Number (n) of protein groups quantified in each condition. Values represent mean ± SD of N = 3. **l** Heat map showing the Pearson correlation of the T47D phosphoproteome shows good reproducibility and variation within experimental conditions. **m** Number (n) of phosphorylated sites in each condition. Values represent mean ± SD of n = 3. **n** Distribution of peptides containing phosphorylated Serine (S), Threonine (T) and Tyrosine (Y) residues (%). **o** Distribution of peptides containing a single, double of more than double modification (%). **p** The distribution of phosphorylated peptides score showed that most of the peptides were identified with high Andromeda score (median: 150.7), consistent with previous[22].

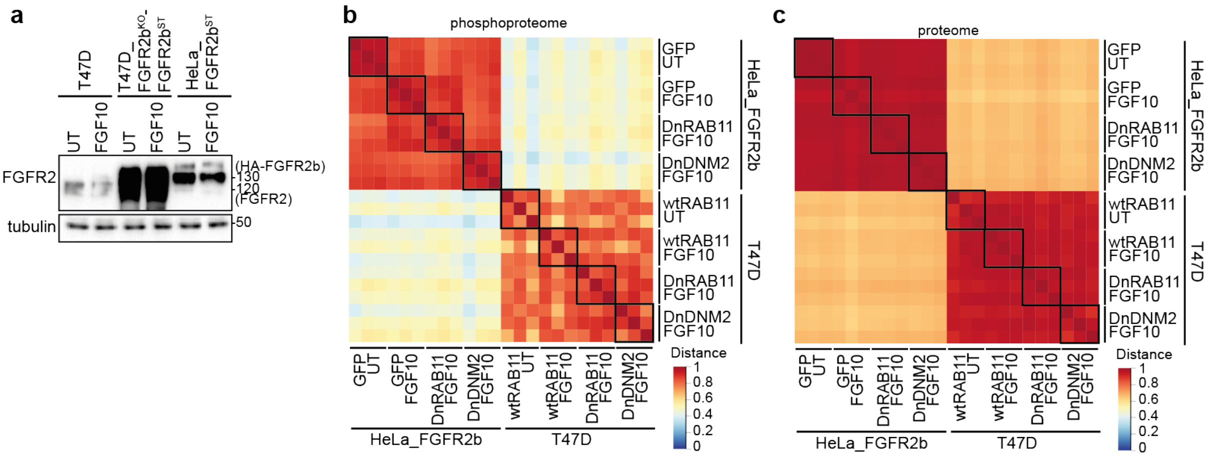


**Supplementary Fig. 2. Quality control of the HeLa and T47D proteomics and phosphoproteomics experiments. a** Immunoblot analysis with the indicated antibodies of parental T47D, T47D_FGFR2b$^{KO}$_FGFR2b$^{ST}$ and HeLa-FGFR2b$^{ST}$ left either untreated (UT) or treated with FGF10 for 40 min. Heat map showing the Pearson correlation of the HeLa vs the T47D proteome (**b**) and phosphoproteome (**c**) shows a significant difference between the two epithelial cell lines.

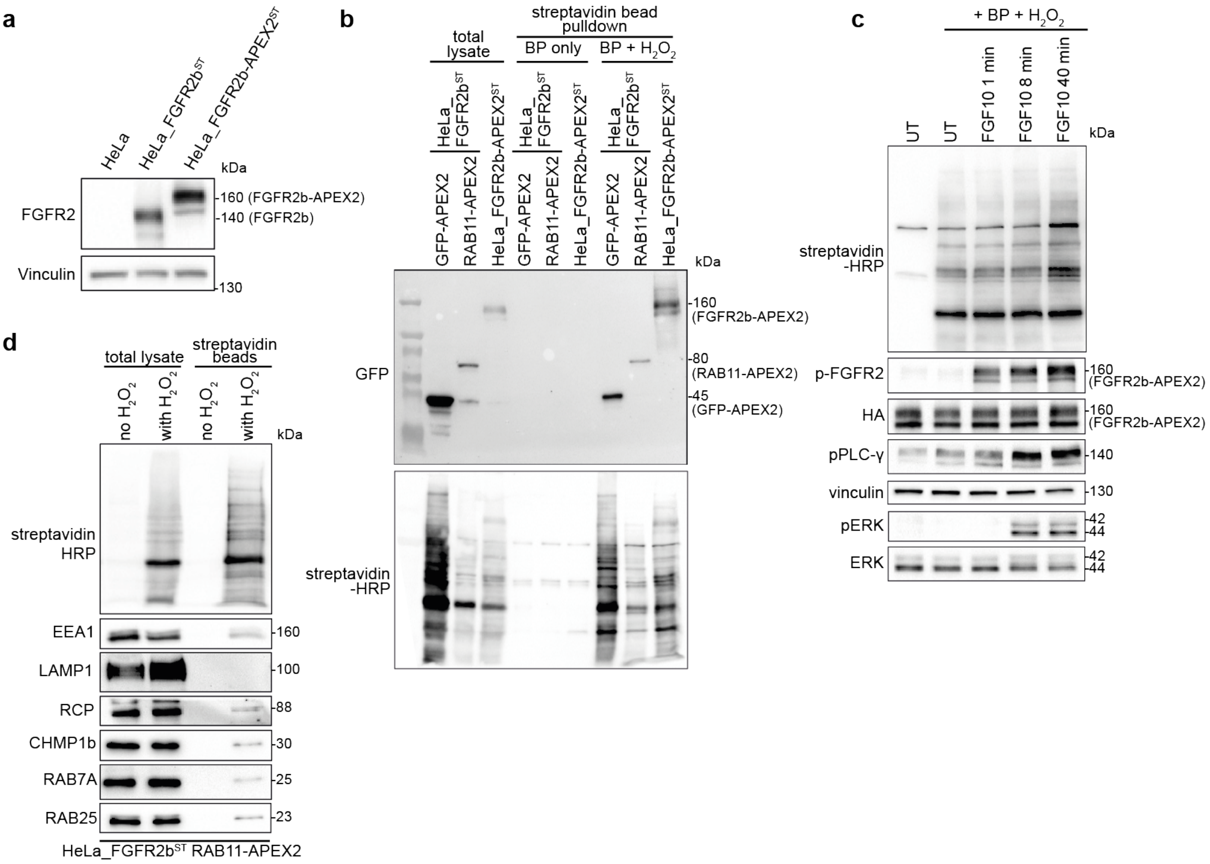


**Supplementary Fig. 3 Quality control of the experiments based on APEX2-tagged proteins. a** Immunoblot analysis with the indicated antibodies of FGFR2b-APEX2 constructs in the indicated cell lines. **b** Immunoblot analysis with the indicated antibodies of HeLa cells transfected with GFP-RAB11-APEX2 or GFP-APEX2 or HA-FGFR2b-APEX2. Total lysates (total) and the pulldown following enrichment of biotinylated samples with streptavidin beads are shown. BP and $H_2O_2$ treatments show that background biotinylation is negligible. **c** Immunoblot analysis with the indicated antibodies of HeLa_FGFR2b-APEX2$^{ST}$ cells stimulated with FGF10 for the indicated time points. **d** Immunoblot analysis with the indicated antibodies of HeLa_FGFR2b$^{ST}$ transfected with RAB11-APEX2.

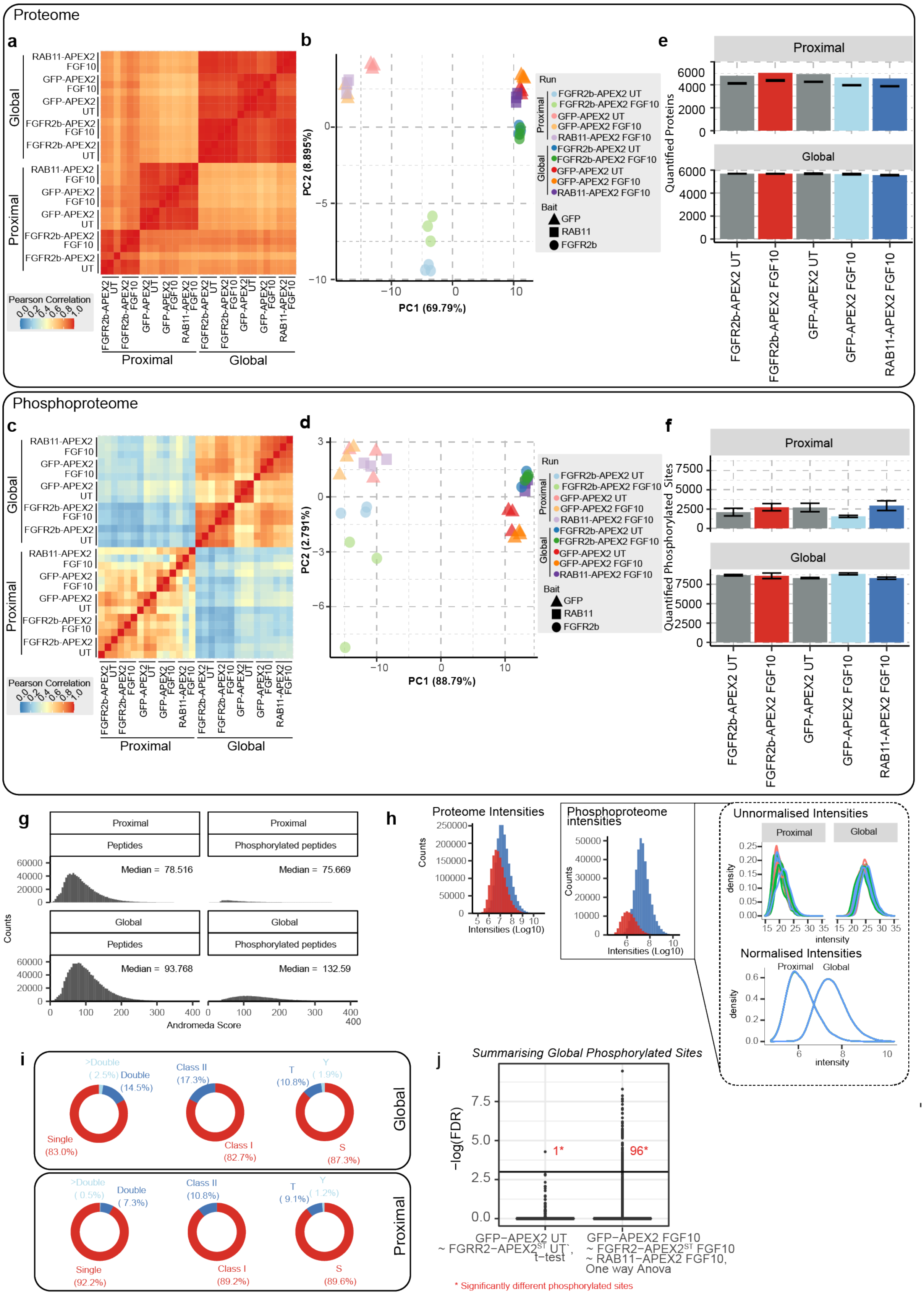


**Supplementary Fig. 4. Quality assessment of proximal and global proteomics and phosphoproteomics. a** Heat map showing the Pearson correlation of the global and proximal proteome of HeLa_FGFR2b-APEX2$^{ST}$ wtRAB11, HeLa_FGFR2$^{ST}$ GFP-APEX2 and HeLa_FGFR2$^{ST}$ RAB11-APEX2 treated with FGF10 or untreated, showed very good reproducibility among biological replicates (Pearson correlation coefficient higher than or equal to 0.9). Differences were seen among UT and FGF10 stimulated cells expressing different APEX2 constructs in the proximal samples, between UT and FGF10 stimulated cells in the global samples, and between the total and proximal samples more broadly (Pearson correlation coefficient smaller than 0.9). **b** PCA of the proteome samples. **c** Heat map showing the Pearson correlation of the phosphoproteome of HeLa_FGFR2b-APEX2$^{ST}$ wtRAB11, HeLa_FGFR2ST GFP-APEX2 and HeLa_FGFR2$^{ST}$ RAB11-APEX2 cells, treated with FGF10 or untreated, showed very good reproducibility among biological replicates (Pearson correlation coefficient higher than or equal to 0.74). Differences were seen among UT and FGF10 stimulated cells expressing different APEX2 constructs in the proximal samples, between UT and FGF10 stimulated cells in the global samples, and between the total and proximal samples more broadly. **d** PCA of the phosphoproteome samples. **e** Number (n) of protein groups quantified in each condition. Values represent mean ± SD of N = 3. **f** Number (n) of phosphorylated sites quantified in each condition. Values represent mean ± SD of n = 3. **g** The distribution of peptides and phosphorylated peptides Andromeda score showed that most of the peptides were identified with relatively high confidence (median score > 75). **h** Distribution of Log10 intensities of the proteome and phosphoproteome from proximal or global experimental conditions. Inset: densities representing the unnormalized and normalized distributions of intensities. Global and proximal proteome samples were normalized together, whilst phosphoproteome samples were normalized separately. See 'Data and Statistical Analysis' section of materials and methods. **i** Visualisation of phosphorylated sites containing a single, double of more than double modification (left); percentage of the phosphorylated sites with a localisation probability ≥ 0.75 (Class I) (%) (middle); and of phosphorylated sites with a modification on Serine (S), Threonine (T) and Tyrosine (Y) residues (%) (right). The two panels represent phosphorylated sites quantified in global (top) or proximal. (bottom) experiments. **j** To assess whether APEX2 tags altered the quantification of the phosphoproteome, statistical tests were performed prior to combining the data from the separate mass spectrometry runs (Fig. 3a and Fig. 4a). A t-test was performed between GFP-APEX2 and FGFR2b-APEX2 UT samples, and a one-way ANOVA between GFP-APEX2, FGFR2b-APEX2 and RAB11-APEX2 treated with 100 ng/ml FGF10 for 40 min. Multiple hypothesis testing was accounted for by calculating the false discovery rate (FDR). In each the UT and FGF10 treated samples, 1 and 96 phosphorylated sites were found to be statistically different due to the APEX2 tags. Given that this was within the 5% chance of statistical error, we concluded that the APEX2 tags did not affect global phosphoproteome quantification and the global UT and FGF10-treated quantitative values could be treated as replicates.

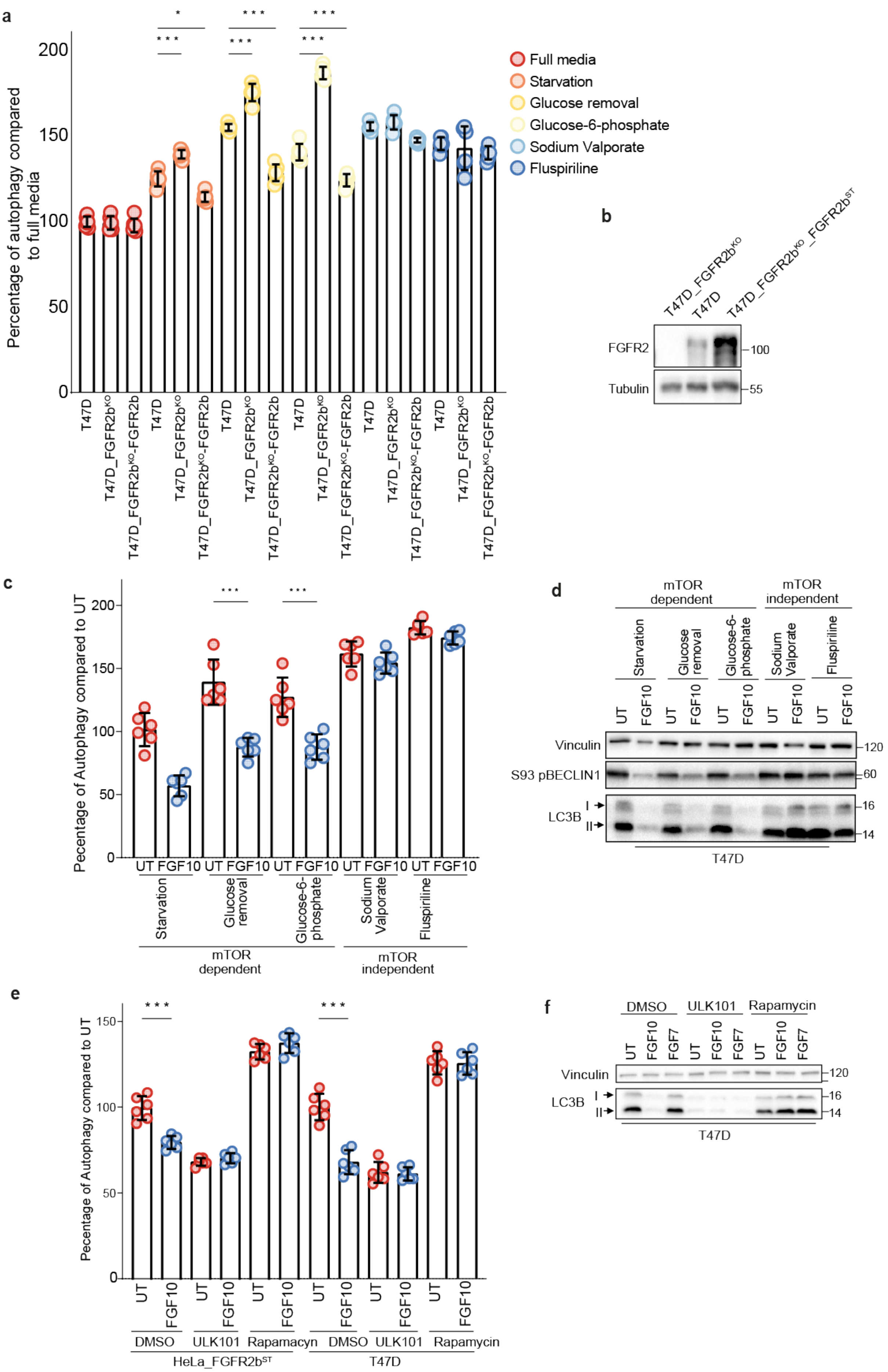
a
Percentage of autophagy compared to full media
0
50
100
150
200
*
***
***
***
***
***
Full media
Starvation
Glucose removal
Glucose-6-phosphate
Sodium Valporate
Fluspiriline
T47D
T47D_FGFR2b$^{KO}$
T47D_FGFR2b$^{KO}$-FGFR2b
b
T47D_FGFR2b$^{KO}$
T47D
T47D_FGFR2b$^{KO}$_FGFR2b$^{ST}$
FGFR2
Tubulin
100
55
c
Pecentage of Autophagy compared to UT
UT FGF10 UT FGF10 UT FGF10 UT FGF10 UT FGF10
Starvation
Glucose removal
Glucose-6-phosphate
Sodium Valporate
Fluspiriline
mTOR dependent
mTOR independent
d
mTOR dependent
mTOR independent
Starvation
Glucose removal
Glucose-6-phosphate
Sodium Valporate
Fluspiriline
UT
FGF10
Vinculin
S93 pBECLIN1
LC3B
I
II
120
60
16
14
T47D
e
Pecentage of Autophagy compared to UT
0
50
100
150
***
***
UT
FGF10
DMSO
ULK101
Rapamacyn
Rapamycin
HeLa_FGFR2b$^{ST}$
T47D
f
DMSO
ULK101
Rapamycin
UT
FGF10
FGF7
Vinculin
LC3B
I
II
120
16
14
T47D

**Supplementary Fig. 5. Role of FGFR2b, mTOR and ULK1 downstream of FGF10 in autophagy regulation. a** Autophagy measured by acridine orange after 24 h treatment in the indicated conditions. N = 6, *P* =< 0.001*** (one-way ANOVA with Tukey test). **b** Immunoblot analysis of FGFR2 expression in T47D, T47D_FGFR2b$^{KO}$ and T47D_FGFR2$^{KO}$_FGFR2b$^{ST}$. **c** Autophagy measured by acridine orange after 2 h treatment of T47D cells in the indicated conditions. N = 6, *P* =< 0.001*** (one-way ANOVA with Tukey test). **d** Immunoblot analysis with the indicated antibodies of T47D treated for 2 h in the indicated conditions. **e** Autophagy measured by acridine orange after 2 h treatment of HeLa_FGFR2b$^{ST}$ and T47D cells in the indicated conditions (See Fig. 7d inset table for inhibitor target). N = 6, *P* =< 0.001*** (one-way ANOVA with Tukey test). **f** Immunoblot analysis with the indicated antibodies of T47D cells treated for 2 h, as indicated.

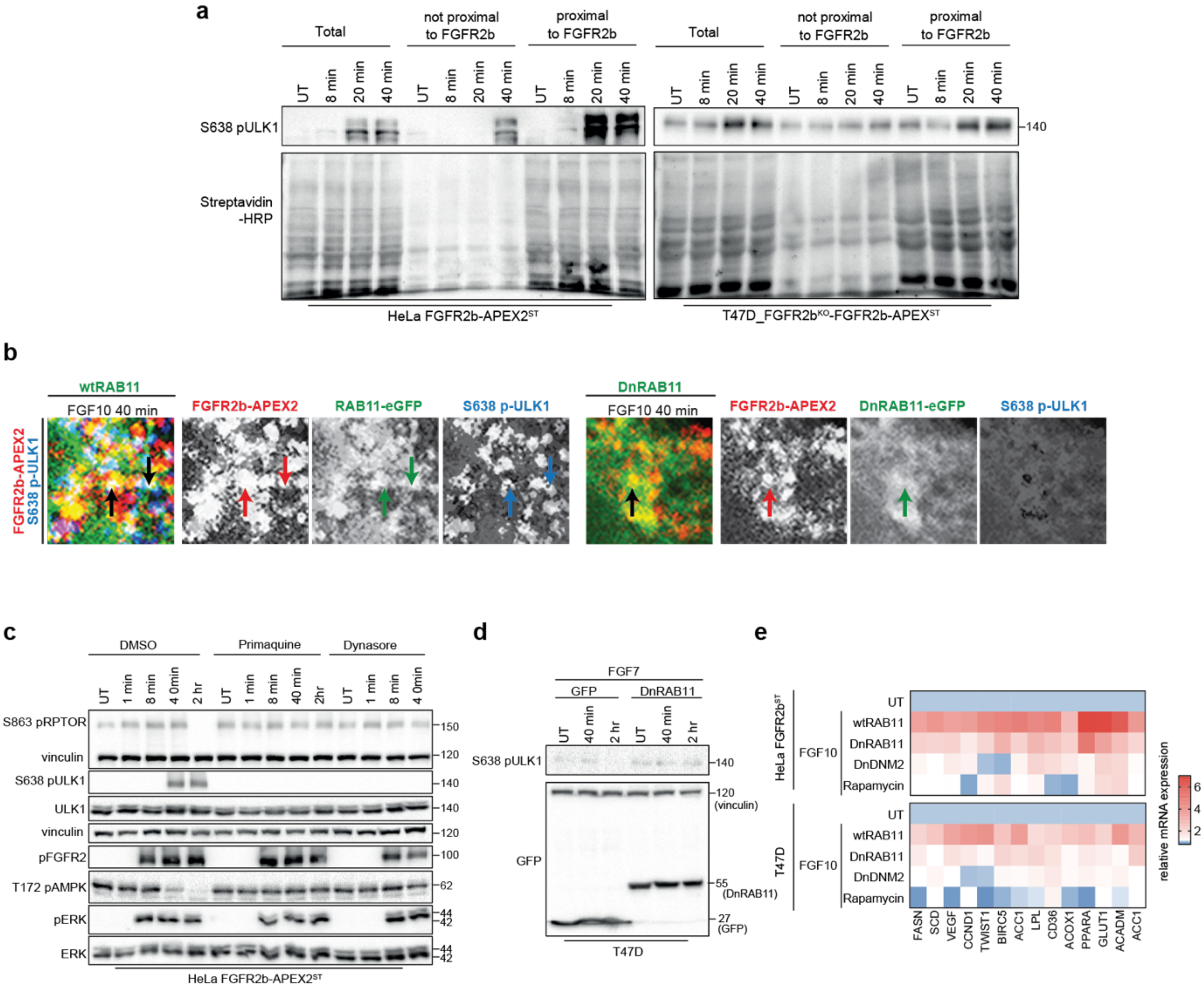


**Supplementary Fig. 6. FGFR2b regulates mTOR and ULK1 signalling from the REs. a** Immunoblot analysis with the indicated antibodies of HeLa FGFR2b-APEX2$^{ST}$ (left) and T47D_FGFR2b$^{KO}$-FGFR2b-APEX2$^{ST}$ (right). Non proximal and proximal samples represent the supernatant and the pulldown following enrichment of biotinylated samples with streptavidin beads, respectively, and run against total lysates (total). **b** Individual panels and merge of the magnified section of FGFR2b-APEX2 (red), phosphorylated ULK1 on S638 (blue) in T47D_FGFR2$^{KO}$_FGFR2b-APEX$^{ST}$ transfected with RAB11 or GFP-DnRAB11 (green) and stimulated or not with FGF10 for 40 min as indicated. Scale bar, 50 µm. Panels from Fig. 6c are shown on the left. **c** Immunoblot analysis with the indicated antibodies of HeLa_FGFR2b$^{ST}$ cells pre-treated with primaquine or Dynasore for 2 h followed by stimulation with FGF10 for the indicated time points. **d** Immunoblot analysis with the indicated antibodies of T47D cells transfected with either GFP or DnRAB11 and stimulated with FGF7 for the indicated time points. **e** Expression of indicated genes in HeLa FGFR2b or T47D transfected with wtRAB11, DnRAB11 or DnDNM2 or pre-incubated with rapamycin for 2 h followed by stimulation with FGF10 for 4 h. qPCR data are presented as heat map from N= 3.

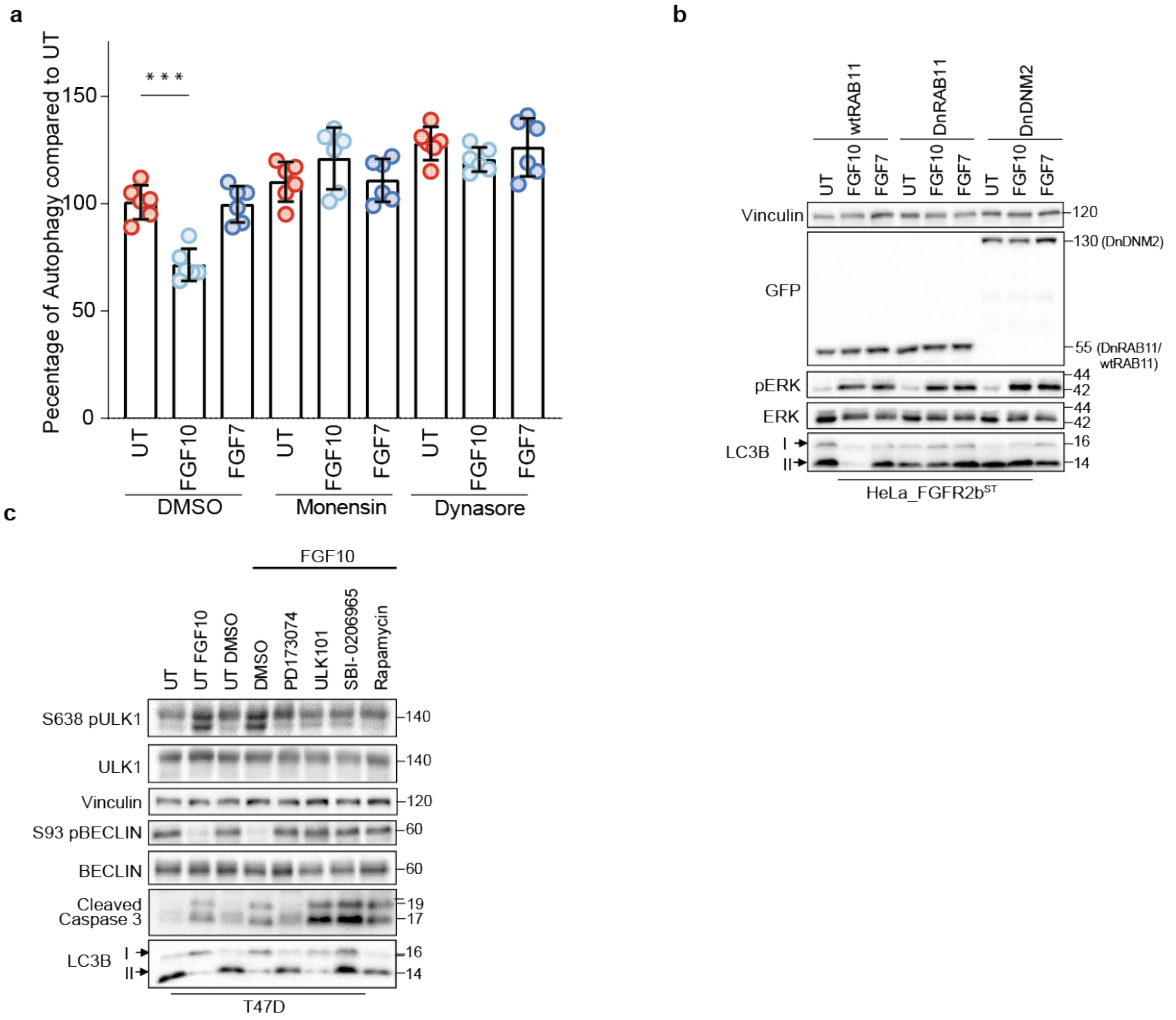


**Supplementary Fig. 7. Inhibiting FGFR2b recycling leads to dysregulated autophagy and an altered balance of proliferation and cell death**. **a** Immunoblot analysis with the indicated antibodies of T47D cells transfected with RAB11 or DnRAB11 or DnDNM2 and stimulated with FGF10 or FGF7 for 2 h. The lysates relate to Fig. 7b. **b** Autophagy measured by acridine orange after treatment of HeLa_FGFR2b$^{ST}$ cells in the indicated conditions. Cells were incubated with monensin or dynasore for 2 h before incubation with FGF7 or FGF10 for 2 h. N = 6, *P* =< 0.001*** (one-way ANOVA with Tukey test **c** Immunoblot analysis with the indicated antibodies of T47D cells pre-treated for 2 h with PD173074, ULK101, SBI0206965, or Rapamycin (targets of inhibition specified in Fig. 7d inset table) and stimulated with FGF10 for 2 h.

**Supplementary Table 1:** HeLa dominant negative normalized, log2 transformed proteome data

**Supplementary Table 2:** HeLa dominant negative normalized, log2 transformed, anova p-value < 0.0001, clustered phosphoproteome data

**Supplementary Table 3:** T47D dominant negative normalized, log2 transformed proteome data

**Supplementary Table 4**: T47D dominant negative normalized, log2 transformed, clustered phosphoproteome data

**Supplementary Table 5:** HeLa APEX2 proteome data

**Supplementary Table 6:** HeLa APEX2 phosphoproteome data
